# Htr6 and Sstr3 ciliary targeting relies on both IC3 loops and C-terminal tails

**DOI:** 10.1101/2020.03.19.997262

**Authors:** Pablo Barbeito, Yuki Tachibana, Raquel Martin-Morales, Paula Moreno, Kirk Mykytyn, Tetsuo Kobayashi, Francesc R. Garcia-Gonzalo

## Abstract

G protein-coupled receptors (GPCRs) are the most common pharmacological target in clinical practice. To perform their signaling functions, many GPCRs must accumulate at primary cilia, microtubule-based plasma membrane protrusions that work as cellular antennae. Despite their great importance, the molecular mechanisms underlying GPCR ciliary targeting remain poorly understood. Serotonin receptor 6 (Htr6) and somatostatin receptor 3 (Sstr3) are two brain-enriched ciliary GPCRs controlling cognition and involved in multiple pathologies such as Alzheimer’s disease and cancer. We previously showed that the third intracellular loops (IC3s) of Htr6 and Sstr3 contain ciliary targeting sequences (CTSs) that are sufficient to confer ciliary localization to non-ciliary GPCRs. However, these CTSs are dispensable for the ciliary targeting of Htr6 and Sstr3 themselves, suggesting these GPCRs have additional CTSs. Herein, we show that the C-terminal tails of Htr6 and Sstr3 also contain CTSs, which act redundantly with those in the IC3s. Accordingly, simultaneous disruption of CTS1 (IC3) and CTS2 (C-terminal tail) abolishes ciliary targeting of both receptors. Mapping the individual residues required for Htr6 ciliary targeting reveals RKQ and LPG motifs critical for CTS1 and CTS2 function, respectively. In Sstr3, CTS1 function relies on the tandem AP[AS]CQ motifs and a subsequent arginine-rich stretch, whereas CTS2 operation requires the juxtamembrane residues. Furthermore, we shed light on the mechanisms of action of Htr6 CTSs by showing how they regulate binding to Tulp3 and Rabl2, two adapters needed for ciliary GPCR targeting.

## INTRODUCTION

Primary cilia are microtubule-based cellular antennae that sense extracellular stimuli in a cell type-specific manner. To do this, each cell type must target specific receptors and signal transducers to its cilia. For instance, limb and neural progenitors in vertebrate embryos must target Patched, Smoothened (Smo) and other Hedgehog (Hh) pathway components to their primary cilia if they are to properly respond to Sonic Hedgehog (Shh), an essential morphogen ^1^. Likewise, kidney epithelial cells must traffic the mechanosensitive polycystin channel to their ciliary membrane in order to sense urine flow ^2^. Failure to do so leads to polycystic kidney disease, the most common of the ciliopathies, a diverse group of human diseases resulting from cilia malfunctions ^3^. In fibroblasts and mesenchymal stem cells, cilia accumulate PDGFRα and IGF1R, two growth factor receptors affecting directional cell migration and adipogenesis, respectively ^4^. Similarly, many neurons throughout the central nervous system (CNS) have primary cilia whose signaling relies on GPCRs and their effectors ^5,6^.

GPCRs are the most common drug target in the human clinic ^7^. The human genome encodes nearly one thousand GPCRs, most of them olfactory receptors (ORs). ORs function in the cilia of olfactory sensory neurons, from nematodes to man. Vision also relies on ciliary GPCR signaling, requiring accumulation of Opsins and their effectors inside retinal photoreceptor cilia. Besides connecting us to the outside world, ciliary GPCRs also sense a myriad of endogenous signals. Some of these GPCRs work outside the nervous system, most in epithelial cells (renal, thyroid, bile duct, airways, endothelium, etc.) but others in mesenchymal stem cells (Ffar4) or fibroblasts, where Smo, Gpr161 and Gpr175 control Hh pathway output. Still, even after discounting visual and olfactory GPCRs, most known ciliary GPCRs function in neurons detecting either neuropeptides (e.g. melanocortin, kisspeptin, galanin, melanin-concentrating hormone, somatostatin) or neuroactive amines like serotonin and dopamine ^5,6,8^.

For olfactory and visual GPCRs, ciliary localization has long been known to be essential for function ^5,6^. This, too, is becoming increasingly clear for other ciliary GPCRs. For instance, some obesity-causing mutations in melanocortin receptor 4 (Mc4r) act by preventing its ciliary targeting, thereby perturbing adenylyl cyclase 3 (AC3)-dependent cAMP signaling in hypothalamic cilia ^9^. Somatostatin receptor 3 (Sstr3) is expressed in cilia throughout the brain, where it also signals via AC3. Remarkably, mice lacking Sstr3, AC3 or cilia in the hippocampus all display very similar learning and memory defects, suggesting that ciliary targeting of Sstr3 and AC3 are essential for these processes ^5^. Sstr3 is also expressed outside the brain, mostly in gastrointestinal tract and testes, and is a promising drug target for diabetes and cancer ^10-12^.

In contrast, serotonin receptor 6 (Htr6) is only expressed in brain, mostly in regions affecting cognition ^13^. Drugs targeting Htr6 hold great promise for treatment of disorders such as anxiety, depression, eating disorders, schizophrenia and Alzheimer’s disease (AD), and as memory enhancers. Although Htr6 also activates Gs-adenylyl cyclase-cAMP signaling, some of its functions are mediated by activation of other effectors such as Cdk5 and the mTOR pathway. Htr6 localizes to neuronal cilia and regulates their length, morphology and composition, effects through which it is reported to affect cognition in a mouse AD model ^13-16^. Thus, it seems very likely that Htr6 ciliary targeting is key for its functions.

The mechanisms underlying ciliary GPCR targeting are only poorly understood. Cis-acting ciliary targeting sequences (CTSs) have been identified in some cases. For some GPCRs, these CTSs are located in their C-terminal tails (CTs). This is the case, among others, of Rhodopsin, Smo and the D1 dopamine receptor ^17-19^). In other instances, however, the CTSs map to the third intracellular loops (IC3s). This was first found for Sstr3 and Htr6 and was later extended to others like Mchr1, Gpr161, Mc4r or Npy2r ^9,20-23^.

For Sstr3 and Htr6, their CTSs were discovered by generating chimeras between these ciliary GPCRs and their non-ciliary relatives Sstr5 and Htr7. After extensive analyses, replacing the IC3s of Sstr5 or Htr7 with those of Sstr3 or Htr6, respectively, was found to suffice for ciliary targeting of the non-ciliary receptors. Furthermore, ciliary targeting of these chimeras was abolished by mutating to phenylalanine the first and last residues of Ax[AS]xQ motifs (hereafter referred to as A-Q motifs) present in the IC3s of both Sstr3 and Htr6 ^20^. In this study, another interesting observation was noted: the reverse pair of chimeras, in which the IC3s of Sstr3 and Htr6 were replaced by those of Sstr5 and Htr7, still accumulated in cilia, indicating that the newly discovered CTSs, albeit sufficient for ciliary targeting of non-ciliary receptors, were dispensable for ciliary targeting of Sstr3 and Htr6 themselves. This suggested, it was also noted, the presence of additional CTSs in these receptors ^20^. A subsequent study confirmed that Htr6 IC3 is dispensable for its ciliary targeting ^16^.

Herein, we report the identification and characterization of those missing CTSs. We show that, for both Sstr3 and Htr6, ciliary targeting occurs as long as either IC3, CT, or both, are present. Conversely, removal of both completely prevents their ciliary accumulation. We then identify the residues required for the function of these CTSs. For Htr6, an LPG motif is critical for C-terminal CTS (CTS2) function, whereas an RKQ motif is key for IC3 CTS (CTS1) function. Interestingly, we also find the A-Q motif in Htr6-IC3 is not needed for CTS1 function and elucidate why mutating A-Q to F-F indeed prevents ciliary targeting of the aforementioned chimera. In contrast, the tandem AP[AS]CQ motifs in Sstr3-IC3 do affect CTS1 function, in conjunction with a neighboring arginine-rich tract. On the other hand, Sstr3 CTS2 function mostly depends on the residues immediately following its seventh transmembrane helix, including LLxP and FK motifs, the latter homologous to the WR motif driving Smo ciliary targeting ^24^.

Finally, we studied how these newly identified CTSs control binding to well-established ciliary trafficking adapters like Tulp3 and Rabl2, both of which interact with the intraflagellar transport (IFT) machinery ^25,26^. We show that Htr6 ciliary targeting is Tulp3-dependent, as previously shown for Sstr3 and other ciliary GPCRs ^22,25,27^. Moreover, we find that the CTs of both Htr6 and Sstr3 associate with Tulp3, in contrast to other Tulp3-dependent GPCRs like Gpr161, whose association is IC3-dependent ^27^. For Htr6, we go on to show that Tulp3 association is mediated by sequences near the LPG motif, which is itself not needed but rather antagonizes Tulp3 association. Thus, Tulp3 dissociation from Htr6 is likely to be an important step for the latter’s ciliary accumulation. Regarding Rabl2, which interacts with Htr6 and is required for its ciliary targeting ^26^, we find that both Htr6 CTSs positively regulate but are not critically required for the Htr6-Rabl2 interaction.

## RESULTS

### Ciliary targeting of Htr6 depends on Tulp3

Many ciliary GPCRs depend on Tulp3 for ciliary targeting. However, whether this is also true for Htr6 has not yet been determined. To clarify this issue, we used HTR6-IMCD3 cells, which we previously generated to stably express Htr6 ^26^. Expression of two independent Tulp3 siRNAs (siTulp3 #1 and #2) in these cells caused a strong reduction in ciliary Htr6 intensity, which was not observed when a negative control luciferase siRNA (siLuc) was expressed instead (Fig.1a-b). Quantitative RT-PCR confirmed that siTulp3 #1 and #2 reduced Tulp3 levels as compared to siLuc (Fig.1c). Thus, Htr6 joins the growing list of ciliary GPCRs shown to depend on Tulp3 for ciliary accumulation ^27^.

**Figure 1.**
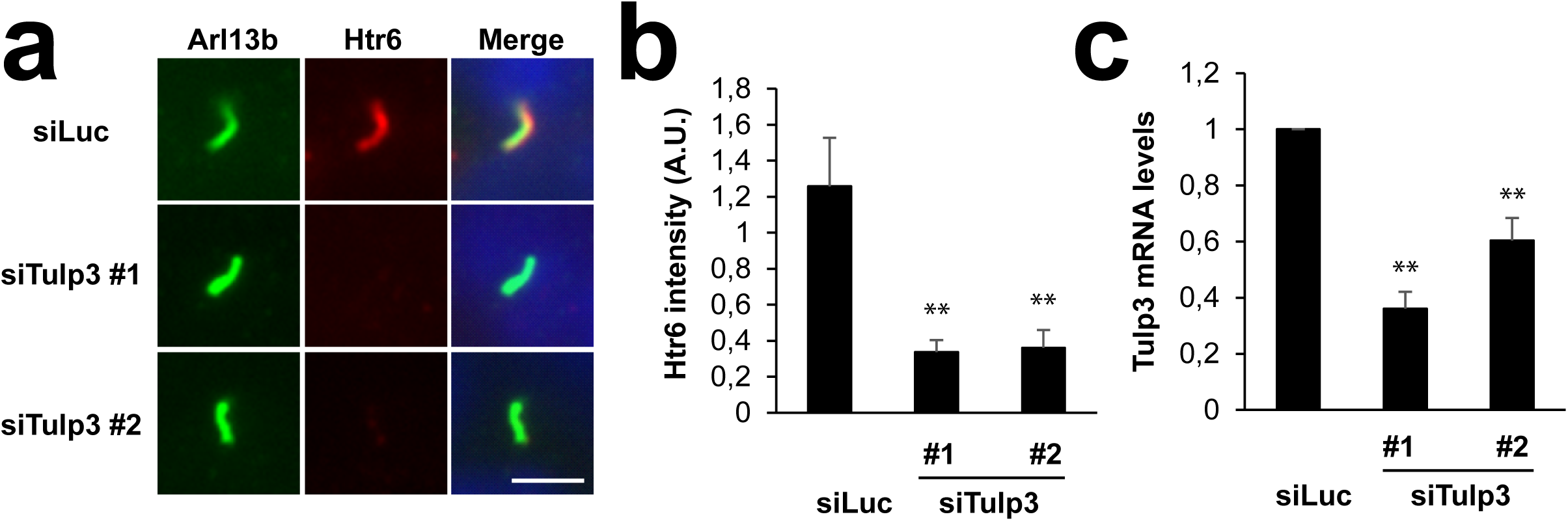
Htr6 ciliary targeting is dependent on the ciliary trafficking adapter Tulp3. **(a)** HTR6-IMCD3 cells transiently transfected with siLuc, siTULP3#1 or siTULP3#2 were cultured in serum-starved medium for 48 hrs and immunostained with anti-Arl13b (green) and anti-Htr6 (red) antibodies. DNA was stained with Hoechst (blue). Scale bar, 2.5 µm. **(b)** Htr6 ciliary intensity was quantified from (a). Data are mean ± SEM of n=23,32,29 cells for siLuc, siTulp3#1 and siTulp3#2, respectively. **(c)** *Tulp3* mRNA levels were analyzed by RT-qPCR and expressed relative to *Gapdh* mRNA. Data are mean±SEM of n=3 independent experiments. Significance in (b-c) is shown as p<0.01(**) in unpaired two-tailed t-tests.

### Htr6-IC3 is sufficient but not necessary for Htr6 ciliary targeting

Htr6 ciliary targeting also involves sequences in its IC3. Indeed, the first half of Htr6-IC3 (aa 208-241) is sufficient to confer cilia localization to Htr7, a non-ciliary GPCR with homology to Htr6, and cilia localization of this chimeric receptor, which we refer to as chimera N, is disrupted by two point mutations (A230F+Q234F) in the ATAGQ motif of Htr6-IC3 ^20^. We confirmed these results (Fig.S1). To determine whether these mutations also disrupt cilia localization of wild type Htr6, we generated a construct expressing Htr6(A230F+Q234F)-EGFP. Interestingly, the resulting protein efficiently accumulates in cilia, to the same extent as Htr6-EGFP wild type control (Fig.S1). This suggests that Htr6-IC3, despite being necessary and sufficient for ciliary targeting of chimera N, is actually dispensable for ciliary targeting of wild type Htr6. Accordingly, Berbari et al. already pointed out in their discussion that Htr6 continues to accumulate in cilia when its Htr6-IC3 is replaced by Htr7-IC3^20^. We confirmed the ciliary targeting of this chimeric receptor, herein referred to as chimera J (Fig.2). Thus, Htr6-IC3 is dispensable for Htr6 ciliary accumulation.

**Figure 2.**
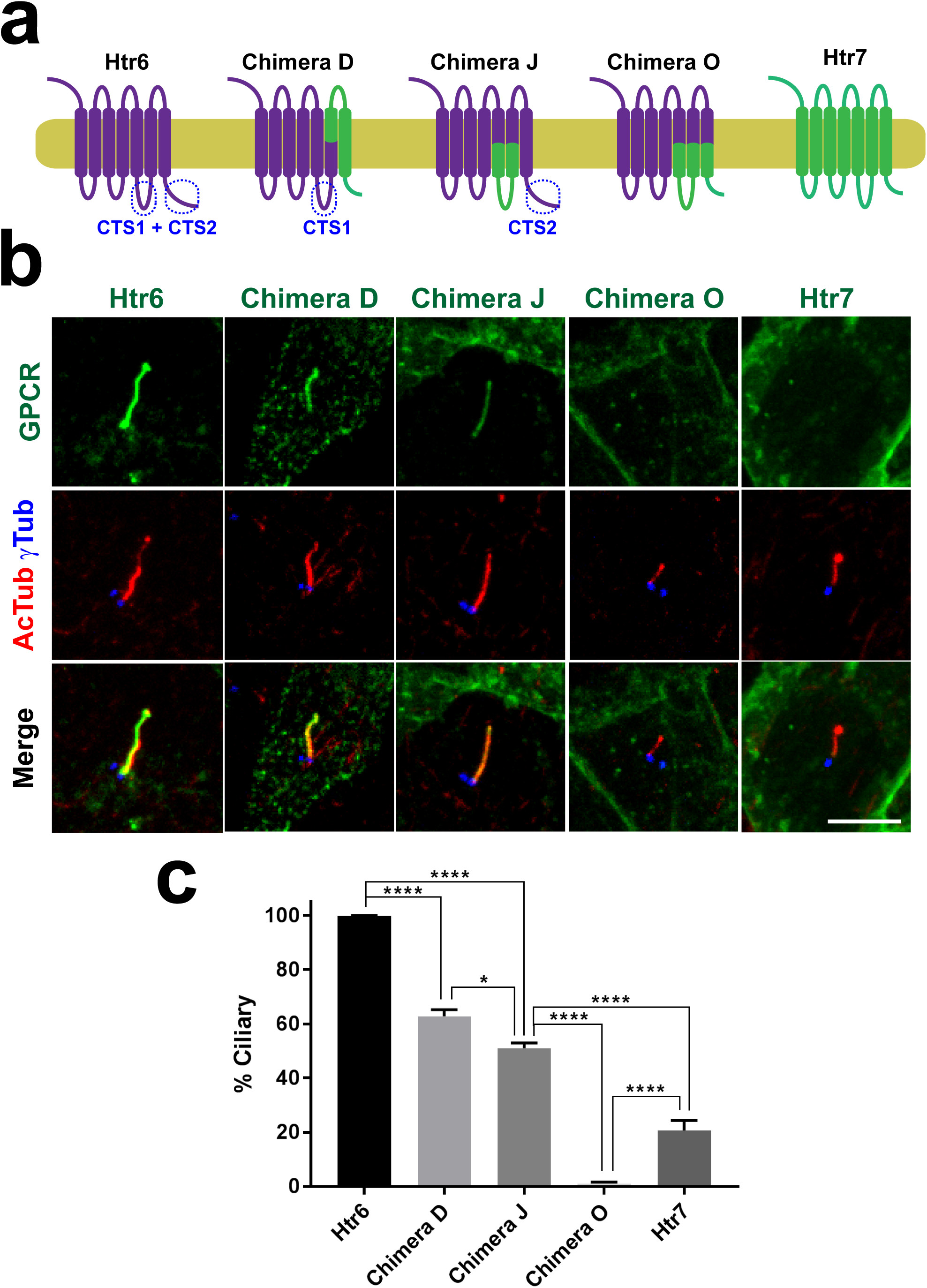
Htr6 ciliary targeting relies on redundant sequences in IC3 loop (CTS1) and C-terminal tail (CTS2). **(a)** Schematic representation of Htr6 (purple), Htr7 (green) and their chimeras, wherein purple segments come from Htr6 and green ones from Htr7. Ciliary targeting sequences in IC3 loop (CTS1) and C-terminal tail (CTS2) are labeled where present. **(b)** The GPCRs from (a), with EGFP fused to their C-termini, were expressed in IMCD3 cells and their cilia localization was analyzed by immunofluorescence with antibodies against EGFP (green), acetylated tubulin (AcTub, red) and gamma-tubulin (γTub, blue). Scale bar, 5 μm. **(c)** Percentage of GPCR-positive cilia in GPCR-transfected cells was quantitated from (b) as described in methods. Data are mean ± SEM of n=5,3,5,5,8 (from left to right) independent experiments, in each of which at least fifty transfected-cell cilia were counted for each GPCR. One-way ANOVA followed by Tukey’s multiple comparisons tests shows all samples are significantly different from each other with p<0.0001 (****) or p<0.05 (*), as indicated.

### Ciliary targeting of Htr6 involves cooperation between IC3 and CT

Since chimera J still accumulates in cilia, we reasoned that Htr6 must contain additional ciliary targeting sequences (CTSs) besides those in its IC3. Since C-terminal tails (CTs) of GPCRs are another common site for CTSs ^17,18^, we tested whether Htr6-CT was necessary for ciliary accumulation of chimera J. To do this, we generated chimera O, which is identical to chimera J except that it contains Htr7-CT instead of Htr6-CT (Fig.2a). When expressed in IMCD3 cells, chimera O completely failed to accumulate in cilia, as opposed to chimera J and wild type Htr6 (Fig.2a-c). Lack of ciliary localization of chimera O was not due to protein instability or retention in Golgi or ER, as chimera O was readily seen at the plasma membrane (Fig.S2). These data indicate that Htr6-CT functions as a CTS in the context of chimera J.

To assess the relative contributions of Htr6-CT and Htr6-IC3 to ciliary targeting of Htr6, we created chimera D, which contains Htr6-IC3 but lacks Htr6-CT (Fig.2a). Like Chimera N (Fig.S1), Chimera D also localized to IMCD3 cilia, confirming that Htr6-CT, like Htr6-IC3, is dispensable for Htr6 ciliary targeting (Fig.2a-b). Altogether, these data indicate that Htr6 ciliary targeting involves redundancy between IC3 and CT: each is sufficient for Htr6 to accumulate in cilia, but none is individually required.

Although chimeras D and J are readily seen in cilia, their ciliary targeting is not as robust as that of wild type Htr6. Upon quantitation, Htr6 robustly localized to virtually all cilia, chimeras D and J were present in about half of them, and chimera O was completely absent from them (Fig.2c). These data seem to indicate that Htr6-IC3 and Htr6-CT are only partially redundant, as each alone is not sufficient for Htr6 cilia localization to be fully penetrant. However, as shown below, this is likely due to chimera use, as Htr6-IC3 and Htr6-CT inactivation by more specific point mutations has no such effects.

### Ciliary targeting function of Htr6-CT maps between residues 392-424

We next set about identifying which amino acid residues constitute Htr6’s CTS2, i.e. the ciliary targeting sequence within Htr6-CT (Fig.2a). Htr6-CT (aa 326-440) is twice as long as Htr7-CT (Fig.3a). Since Htr6-CT’s first half can be aligned with Htr7-CT, we generated another chimera by swapping the first half of Htr6-CT in chimera J by Htr7-CT. The resulting protein, chimera Q, still localized to cilia, indicating that aa 326-368 of Htr6-CT are not essential for CTS2 function (Fig.3a-d). Consistent with this, ciliary targeting of Htr6 was not affected by two mutations, S352A and S352D, which substitute non-phosphorylatable (Ala) and phosphomimetic (Asp) residues for Ser352, the mouse residue whose counterpart in human HTR6 is phosphorylated by Cdk5 kinase (Fig.S3) ^28^. Thus, the critical CTS2 residues should localize within Htr6-CT’s second half. Accordingly, deletion of Htr6-CT’s second half (Δ373-440) or last third (Δ401-440) completely abolishes cilia localization of chimera J (Fig.3a-d), even though these mutants have no problem reaching the plasma membrane (Fig.S2). In contrast, deletions Δ369-370, Δ371-378, Δ379-391 and Δ425-440 do not abolish ciliary targeting, even if Δ425-440, like chimera Q, moderately reduces it (Fig.3a-d). Altogether, these data indicate that CTS2 function critically requires residues within aa 392-424, and that some residues outside this critical region reinforce CTS2 action.

**Figure 3.**
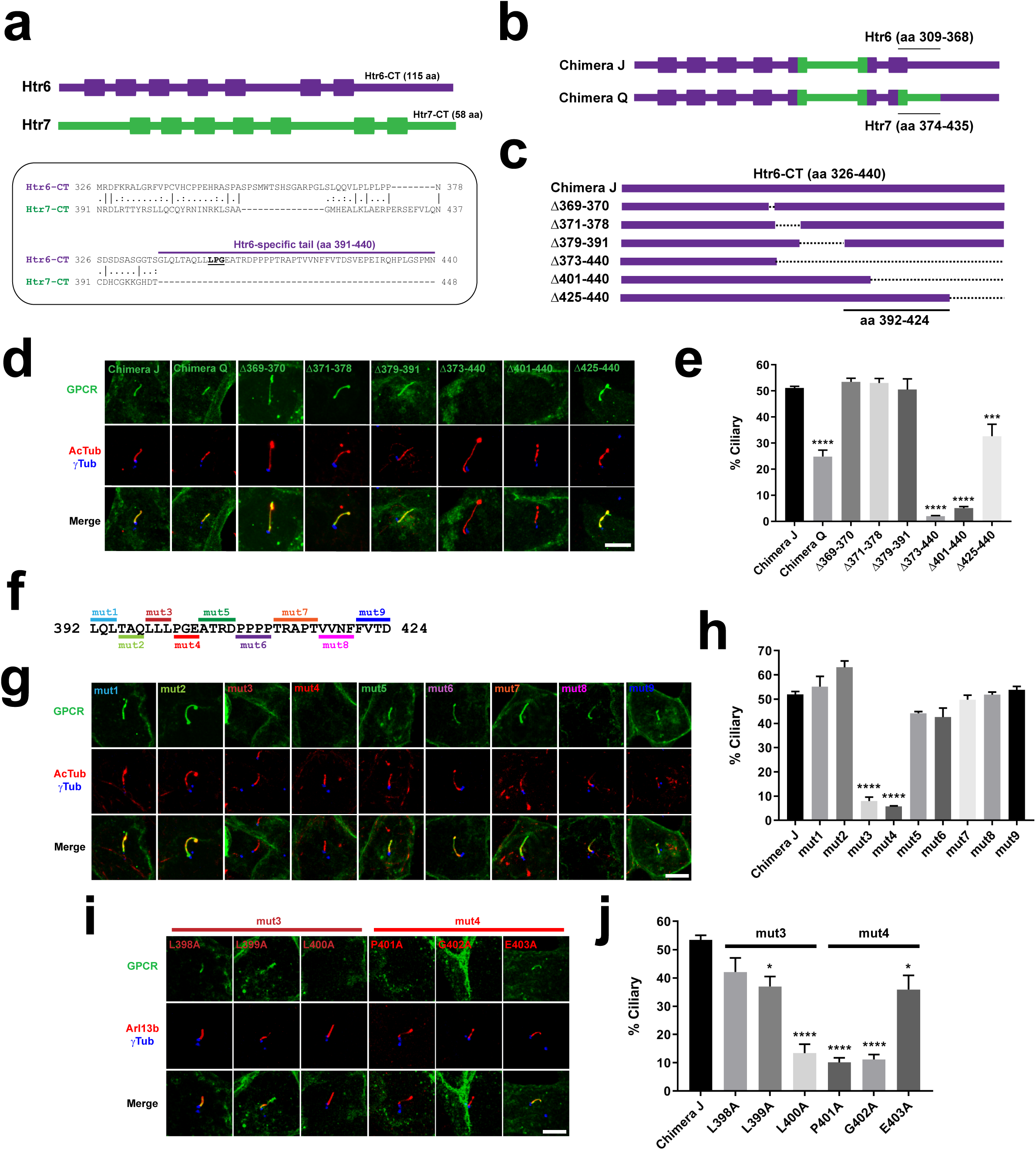
An LPG motif is critical for the ciliary targeting function of Htr6’s CTS2. **(a)** Top: schematic depiction of Htr6 (440 aa, purple) and Htr7 (448 aa, green) with transmembrane helices displayed as boxes. Notice how Htr6 C-terminal tail (CT) is twice as long as Htr7-CT (115 aa versus 58 aa). Bottom: alignment of Htr6-CT and Htr7-CT. The former is 10-fold richer in prolines (17% vs. 1.7%) and its latter half (aa 391-440) has no homologous counterpart in Htr7. LPG motif inside Htr6-specific tail is underlined. **(b)** Schematic representation of Chimera J and Chimera Q. They are identical except that Chimera Q lacks the Htr6 residues indicated in Chimera J, and contains instead the Htr7 residues indicated in Chimera Q. **(c)** Schematic showing Htr6-CT on top (present in Chimera J), and the deletions that were introduced into Chimera J, covering all residues that had not been substituted in Chimera Q. Indicated at the bottom is the critical region required for ciliary targeting of Chimera J. **(d)** IMCD3 cells expressing the constructs from (b-c), all fused to EGFP in their C-termini, were analyzed by immunofluorescence with antibodies against EGFP (green), acetylated tubulin (AcTub, red) and gamma-tubulin (γTub, blue). Scale bar, 5 μm. **(e)** Percentage of GPCR-positive cilia relative to total transfected-cell cilia was quantitated from (d). **(f)** Nine separate mutations (mut1-mut9) were introduced into critical region from (c), with all indicated residues replaced by alanines. **(g)** IMCD3 cells expressing C-terminally EGFP-tagged mut1-mut9 chimera J mutants were analyzed as in (d). Scale bar, 5 μm. **(h)** Percentage of GPCR-positive cilia from (g) was quantitated as in (e). **(i)** The residues mutated in mut3 and mut4 were individually substituted to alanine and analyzed as in (d). **(j)** Percentage of GPCR-positive cilia from (i) was quantitated as in (e). In all quantitations, data are mean ± SEM of n=3-5 independent experiments per construct, with at least fifty cilia counted per construct and experiment. Statistical analysis was performed by one-way ANOVA followed by Tukey’s multiple comparisons tests. Significance is indicated as p<0.05(*), p<0.001(***) or p<0.0001(****).

### Ciliary targeting function of Htr6-CT is mediated by an LPG motif

To pinpoint which residues inside the critical region are required for CTS2 function, we generated nine alanine-scanning mutants (mut1-mut9), together spanning all 33 residues in the region (Fig.3f). Seven of these mutants had no effect on cilia localization of chimera J, while the other two mutants abolished it (Fig.3f-h). These two mutants, mut3 (LLL398-400AAA) and mut4 (PGE401-403AAA), still reached the plasma membrane, indicating that their absence from cilia is not due to lack of expression or failure to exit ER or Golgi (Fig.S2). Next, we individually mutated to alanine each of the six residues covered by mut3 and mut4. Mutants L398A, L399A and E403A did not significantly reduce ciliary targeting of chimera J, while L400A, P401A and G402A clearly did (Fig.3i-j). Thus, L400, P401 and G402 are the three key residues for Htr6 CTS2 function.

### Ciliary targeting function of Htr6-IC3 is mediated by an RKQ motif

Aside from the A230F+Q234F mutation in Htr6-IC3 interfering with chimera N ciliary targeting ^20^ (Fig.S1), nothing is known about the exact residues mediating CTS1 function in Htr6-IC3. To clarify this, we first introduced the mut3 mutation from Fig.3 (henceforth CT-mut3) into wild type Htr6 and checked its cilia localization, which was indistinguishable from wild type Htr6 (Fig.4). Since CT-mut3 abolishes CTS2 function (Fig.3g-h), ciliary targeting of Htr6(CT-mut3) must depend on CTS1 function. Thus, we combined CT-mut3 with IC3 mutations in order to map CTS1 function (Fig.4a-b). Since Htr6-IC3 residues 208-241 are sufficient for ciliary targeting of chimera N (Fig.S1) ^20^, we started by making three deletions spanning this sequence (Δ208-219, Δ220-229 and Δ230-241) and combining them with CT-mut3 (Fig.4a-d). The first two deletions (IC3-Δ1 and IC3-Δ2) abolished ciliary targeting of Htr6(CT-mut3), whereas the last deletion (IC3-Δ3) had no effect (Fig.4c-d). The strongest effect was seen for Htr6(IC3-Δ1+CT-mut3). Although this protein reaches the plasma membrane, it does so less efficiently, consistent with Δ208-219 disrupting a tyrosine-based sorting motif (208-YxxI-211) right after Htr6’s fifth transmembrane helix (Fig.4a, Fig.S2) ^29^. For Htr6(IC3-Δ2+CT-mut3) the loss of ciliary targeting was strong but not complete, and this protein readily reached the plasma membrane (Fig.4c-d, Fig.S2). Thus, residues 208-229, but not 230-241, are important for CTS1 function.

**Figure 4.**
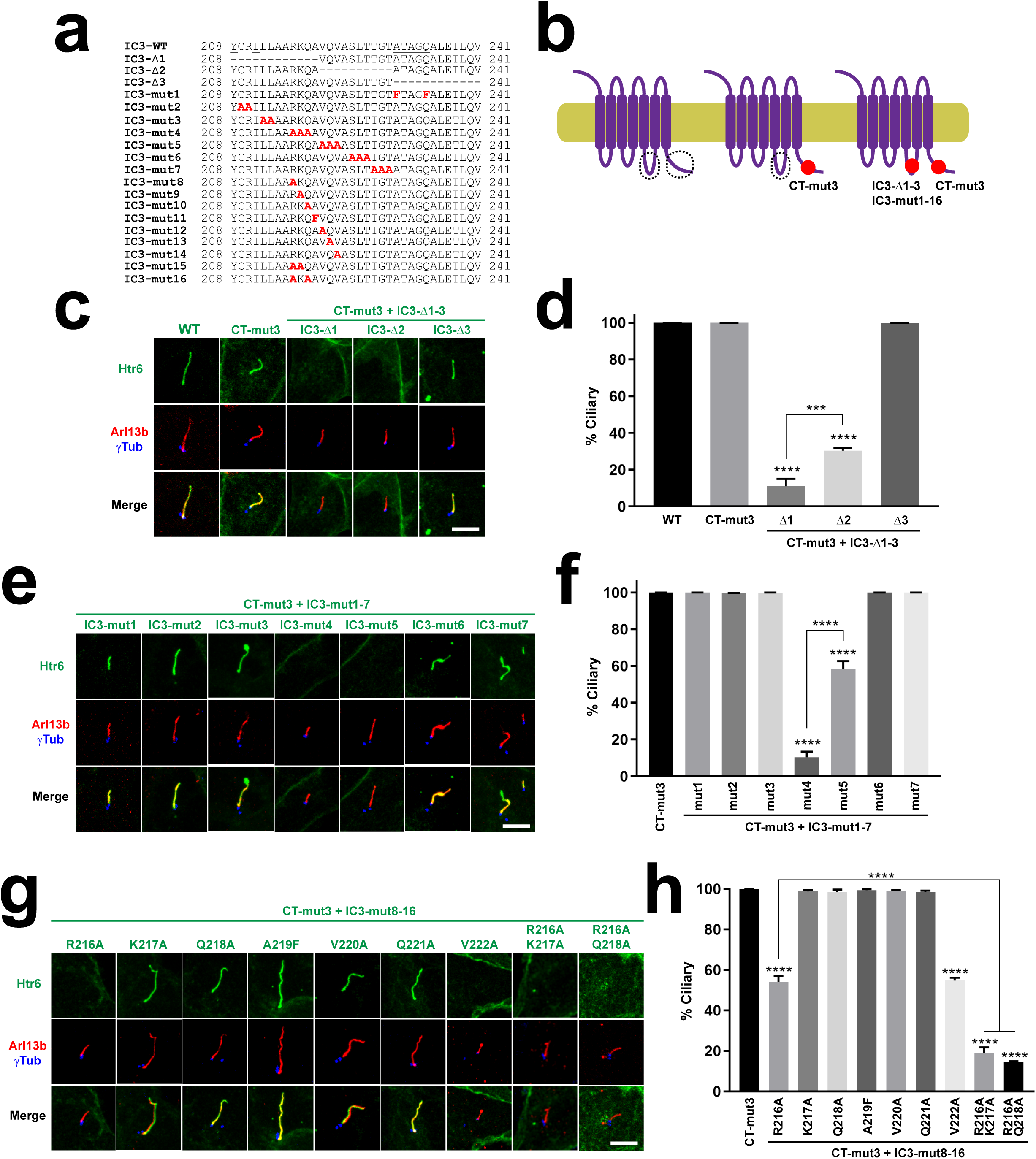
An RKQ motif is critical for IC3-dependent Htr6 ciliary targeting. **(a)** Sequence of Htr6’s IC3 loop and its mutants used here. **(b)** Schematic of Htr6 wild type and its mutants used here. CT-mut3 is the mut3 CTS2 mutation from Figure 3. The IC3 mutations from (a) were combined with CT-mut3. Mutations shown as red spots. CTS1 and CTS2 encircled with dashed lines when intact. **(c)** IMCD3 cells expressing the indicated versions of Htr6, all fused to C-terminal EGFP, were analyzed by immunofluorescence with antibodies against EGFP (green), Arl13b (red) and gamma-tubulin (γTub, blue). Scale bar, 5 μm. **(d)** Percentage of GPCR-positive cilia relative to total transfected-cell cilia was quantitated for the constructs in (c). **(e)** The indicated Htr6 mutants were analyzed as in (c). **(f)** Ciliary targeting of Htr6 mutants from (e) was quantitated as in (d). **(g)** The indicated Htr6 mutants were analyzed as in (c). **(h)** Ciliary targeting of Htr6 mutants from (g) was quantitated as in (d). Data in (d), (f) and (h) are mean ± SEM of n=3-4 independent experiments per construct, with at least fifty cilia counted per construct and experiment. Data were analyzed by one-way ANOVA followed by Tukey’s multiple comparisons tests. Unless otherwise indicated, significance is shown relative to control sample (black column) with p<0.001(***) or p<0.0001(****).

The lack of effect by IC3-Δ3 was surprising, as it deletes the A-Q motif (230-ATAGQ-234), whose A230F+Q234F mutation (IC3-mut1) disrupts chimera N ciliary targeting (Fig.S1) ^20^. Since IC3-mut1 introduces two bulky phenylalanine residues in place of smaller A230 and Q234, we hypothesized that IC3-mut1 effect on chimera N might be due to dominant negative effects on CTS1 function, rather than on A230 and Q234 being needed. If so, then Htr6(IC3-mut1+CT-mut3) should also fail to localize to cilia. However, its cilia localization is as good as that of wild type Htr6 (Fig.4e-f). Thus, the A-Q to F-F mutation does not interfere with CTS1 function of Htr6. And yet, it somehow prevents chimera N from reaching cilia (Fig.S1a-c). Upon careful analysis of the A-Q and F-F versions of chimera N, we discovered that the F-F version fails to reach the plasma membrane ≈7-fold more often than the A-Q version. This likely explains why the F-F version’s cilia localization is ≈5-fold lower than the A-Q version’s (Fig.S1d-e).

To refine CTS1 mapping, we next created six alanine-scanning mutations (IC3-mut2 to IC3-mut7) spanning residues 208-229, with the exception of Y208 and I211 to avoid disrupting the aforementioned sorting motif (Fig.4a). Of these six mutants, four localized to cilia and two failed to do so (Fig.4e-f). These two mutants, IC3-mut4 (RKQ216-218AAA) and IC3-mut5 (VQV220-222AAA), were abundantly seen at the plasma membrane, showing that their effect on ciliary targeting is specific (Fig.S2). Quantitatively, IC3-mut4 had a much stronger effect than IC3-mut5 (10-fold versus 1.7-fold reduction), indicating that 216-RKQ-218 are key for CTS1 function. As done above for CTS2, we individually mutated each residue in IC3-mut4 and IC3-mut5 to alanine. We also created the A219F mutant, so that the entire 216-RKQAVQV-222 stretch was covered. Analysis of IMCD3 cilia localization of these seven mutants showed that R216A and V222A reduce in half the ciliary targeting of Htr6(CT-mut3), while the other mutants do not affect it (Fig.4g-h). Thus, V222 fully accounts for the effect seen with IC3-mut5, whereas R216 only partially accounts for the stronger reduction seem with IC3-mut4. This suggests that K217 and/or Q218 positively contribute to CTS1 activity when R216 is absent. To test for this, we created the R216A+K217A and R216A+Q218A mutants, both of which phenocopy IC3-mut4 (Fig.4g-h). Thus, Htr6 CTS1 function critically depends on the RKQ triad, within which R216 is the most important residue while K217 and Q218 play ancillary roles.

Intriguingly, the RKQ triad is preceded by two alanines, making it a non-canonical A-Q motif (canonical being Ax[AS]xQ ^20^). However, whether alanines 214-215 play a role in Htr6 CTS1 function remains to be explored.

### Htr6 CT and IC3 are both sufficient for ciliary targeting

Thus far, our data indicate that Htr6 cilia localization is mediated by cooperation between two redundant ciliary targeting sequences, CTS1 and CTS2, located in IC3 and CT, respectively. Therefore, in the context of Htr6-Htr7 chimeras or of Htr6 mutants, both CTS1 and CTS2 are sufficient to drive ciliary targeting. For CTS2, we further confirmed this by fusing Htr6-CT at the C-terminal end of Htr7 (Fig.5a). The resulting Htr7-(Htr6-CT) fusion protein strongly accumulates in cilia, showing that Htr6-CT is sufficient to target Htr7 to cilia (Fig.5b-c). We then tested whether Htr6-CT also suffices to target a single-pass transmembrane protein to cilia. To do this, we substituted Htr6-CT for the cytosolic domain of CD8α, a single transmembrane protein that has repeatedly been used for this same purpose ^27,30^. Indeed, the CD8α-(Htr6-CT) chimera readily accumulated in cilia, as did CD8α-(Htr6-IC3). By contrast, CD8α-(Htr7-CT) failed to accumulate in cilia (Fig.5d-f), indicating that both Htr6-CT and Htr6-IC3 are sufficient to specifically target a single transmembrane protein to cilia. Lastly, we also checked whether Htr6-CT is sufficient to target a soluble protein to cilia. To test this, we created the (Htr6-CT)-EGFP fusion protein, which failed to accumulate in cilia, indicating that CTS2 function of Htr6-CT requires membrane association (Fig.S4).

**Figure 5.**
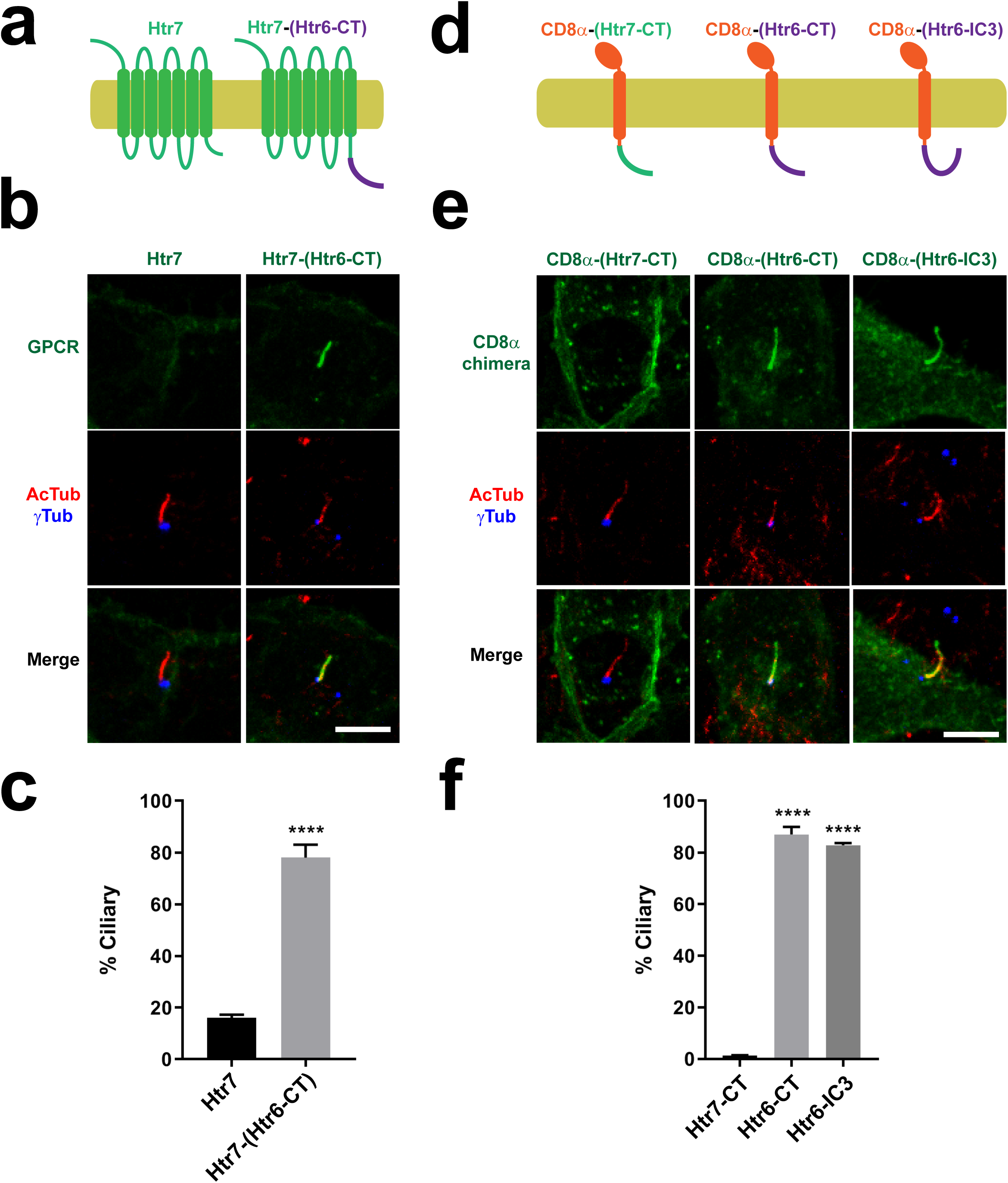
Htr6 CTS1 and CTS2 are both sufficient for ciliary targeting. **(a)** Schematic of Htr7 with or without the CTS2-containing C-terminal tail of Htr6 (Htr6-CT) fused to its C-terminus. **(b)** IMCD3 cells expressing C-terminally EGFP-tagged constructs from (a) were analyzed by immunofluorescence with antibodies against EGFP (green), acetylated tubulin (AcTub, red) and gamma-tubulin (γTub, blue). Scale bar, 5 μm. **(c)** Percentage of GPCR-positive cilia relative to total transfected-cell cilia was quantitated from (b). **(d)** Schematic of CD8α(1-206) chimeras, containing extracellular and transmembrane regions of CD8α fused to Htr7-CT, Htr6-CT (containing CTS2) or Htr6-IC3 (containing CTS1). **(e)** IMCD3 cells expressing C-terminally EYFP-tagged constructs from (d) were analyzed as in (b). Scale bar, 5 μm. **(f)** Percentage of GPCR-positive cilia relative to total transfected-cell cilia was quantitated from (e). Data in (c) and (f) are mean ± SEM of n=4-5 independent experiments per construct. In each experiment, at least 50 cilia were counted per condition. Data were analyzed by unpaired two-tailed t-test (c) or by one-way ANOVA followed by Tukey’s multiple comparisons tests (f). Significance in both cases is shown as p<0.0001 (****).

### Sstr3 ciliary targeting also involves redundant CTSs at IC3 and CT

In their study, Berbari et al. showed that Htr6 ciliary targeting mechanisms resemble those of Sstr3^20^. Furthermore, Sstr3-IC3 was also seen to be sufficient but not necessary for cilia localization, as replacement of Sstr3-IC3 by the IC3 of non-ciliary Sstr5 did not abolish ciliary targeting ^20^. This observation, puzzling at the time, may now readily be explained if Sstr3-CT also contains a CTS2. To test this, we first checked whether Sstr3-CT is sufficient to drive cilia localization of a CD8α-(Sstr3-CT) chimera. Indeed, both CD8α-(Sstr3-CT) and CD8α-(Sstr3-IC3) chimeras accumulate in cilia, with the former doing so even more strongly than the latter (Fig.6a-c). Therefore, both Sstr3-IC3 and Sstr3-CT contain ciliary targeting sequences.

**Figure 6.**
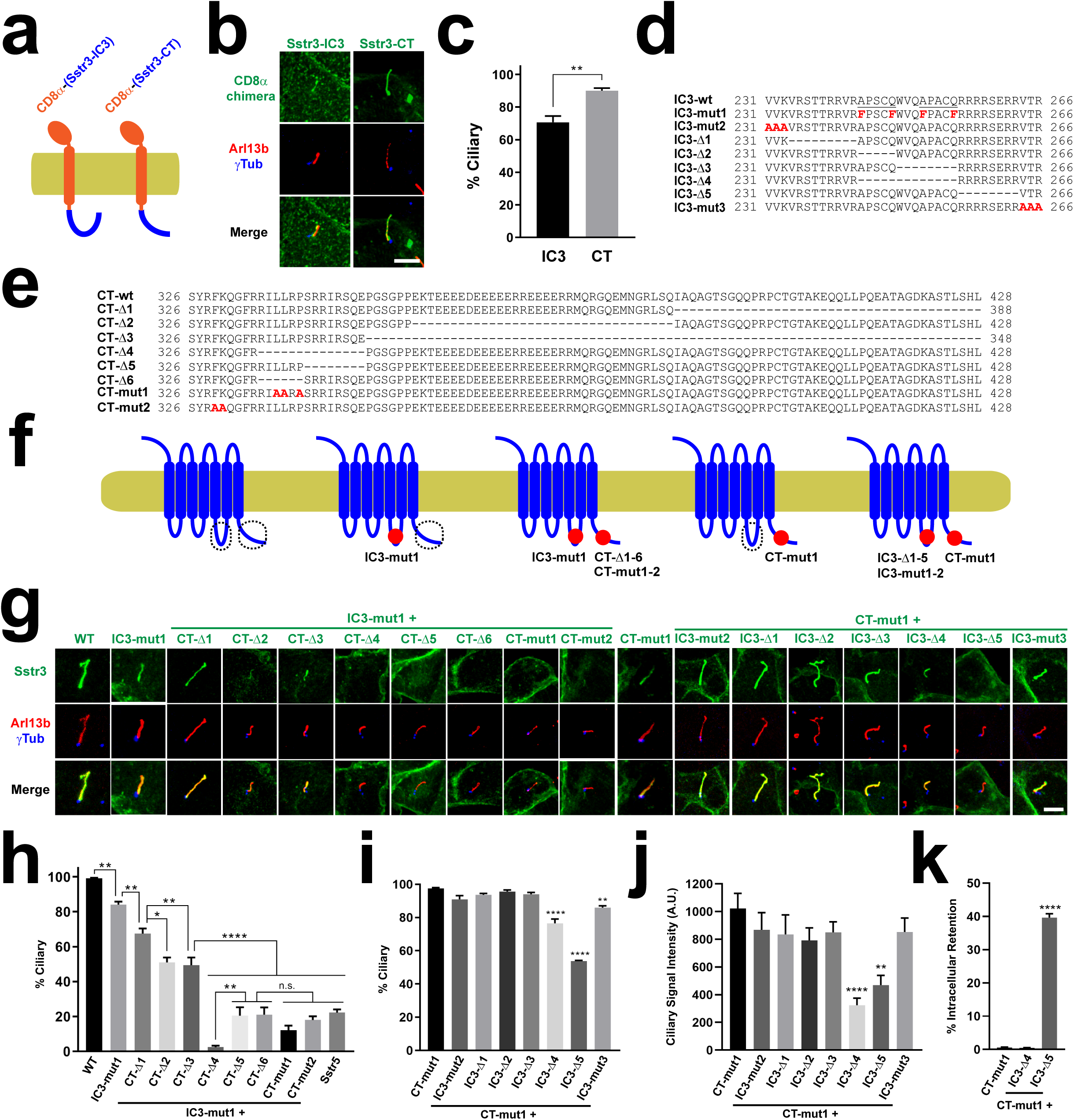
Sstr3 ciliary targeting also depends on redundant ciliary targeting sequences in IC3 and CT. **(a)** Schematic of CD8α(1-206) chimeras fused to Sstr3-IC3 or Sstr3-CT. **(b)** IMCD3 cells expressing C-terminally EYFP-tagged constructs from (a) were analyzed by immunofluorescence with antibodies against EGFP/EYFP (green), Arl13b (red), and gamma-tubulin (γTub, blue). Scale bar, 5 μm. **(c)** Percentage of GPCR-positive cilia relative to total transfected-cell cilia was quantitated for the constructs in (b). Data are mean ± SEM of n=4,8 (IC3,CT) independent experiments per construct. Significance in unpaired two-tailed t-test shown as p<0.01(**). **(d)** Sstr3-IC3 wild type sequence (top) and its mutated versions used below. The two reported Ax(A/S)xQ motifs are underlined in wild type sequence. **(e)** Sstr3-CT wild type sequence (top) and its mutated versions used below. **(f)** Schematic of Sstr3 and its mutants used below. CTS1 and CTS2 are encircled where intact. Mutations shown as red spots. **(g)** Ciliary targeting of Sstr3 mutants from (f) was analyzed as in (b). **(h-i)** Percentage of positive cilia for each of the indicated Sstr3 constructs from (g) was quantitated as in (c). **(j)** Intensity of ciliary staining was quantitated for the indicated Sstr3 constructs. **(k)** Percentage of cells with no detectable plasma membrane or ciliary staining was quantitated for indicated constructs. Data in (h-k) are mean ± SEM and were analyzed by one-way ANOVA followed by Tukey’s multiple comparisons tests. Significance shown as p<0.05(*), p<0.01(**), p<0.001(***), p<0.0001(****) or not significant (n.s.). For (h), numbers of independent experiments per construct from left to right were n=10,3,4,3,3,3,4,4,3,4,10. Equivalent numbers for (i) were n=3,4,4,5,5,5,4,4. For both (h) and (i), at least 50 cilia were counted per construct and experiment. For (j), intensity was measured in n=26-59 cilia per condition in one representative experiment. For (k), n=4 independent experiments per construct with at least 200 transfected cells assessed per construct and experiment.

### Ciliary targeting function of Sstr3-CT is mediated by its juxtamembrane region

As done with Htr6, we then characterized how the two CTSs functionally relate to one another and which residues underlie their function. For this, we generated a series of deletion and point mutants in both Sstr3-IC3 and Sstr3-CT (Fig.6d-f). Previously, a mutant was already identified in Sstr3-IC3 that disrupts ciliary targeting of an Sstr3-Sstr5 chimera containing Sstr3-IC3 in an Sstr5 background^20^. When this mutation (A243F+Q247F+A251F+Q255F, henceforth IC3-mut1, Fig.6d) was introduced to wild type Sstr3, cilia localization was only very mildly affected, reducing the percentage of Sstr3-positive cilia from ≈100% to ≈80% (Fig.6g-h). We then examined whether ciliary targeting of Sstr3(IC3-mut1) is disrupted by mutations in Sstr3-CT (aa 326-428) (Fig.6e-f).

Deleting the last third of Sstr3-CT (CT-Δ1: Δ389-428) from Sstr3(IC3-mut1) caused another mild reduction in ciliary targeting, from ≈80% to ≈65% (Fig.6g-h). Deleting the central third of Sstr3-CT (CT-Δ2: Δ355-388), which contains a coiled coil highly enriched in glutamate residues, had a stronger effect, reducing targeting from ≈80% to ≈50% (Fig.6g-h). A very similar effect was observed with CT-Δ3 (Δ349-428), which gets rid of all but the first 23 aa of Sstr3-CT (Fig.6g-h). Thus, although Sstr3-CT aa 349-428 modulate ciliary targeting, they are not critical for it.

A mutation deleting aa 335-428 did prevent ciliary but also plasma membrane targeting, and was accumulated intracellularly (Fig.S5). Instead, internal deletion of aa 335-348 (CT-Δ4) completely abolished ciliary targeting without affecting trafficking to plasma membrane (Fig.6f-g, Fig.S5). Deleting either the first (CT-Δ5: Δ335-RILLRP-340) or second (CT-Δ6: Δ341-SRRIRSQE-348) half of this sequence also abolished ciliary targeting, leaving only ≈20% of mildly positive cilia, the same as in the non-ciliary Sstr5 negative control (Fig.6g-h) ^20^. None of these mutations obstructed plasma membrane trafficking (Fig.S5). Hence, the region 335-348 contains essential residues for CTS2 function of Sstr3. Among these residues there is an LLxP motif reminiscent of Rhodopsin’s VxP and other ciliary targeting sequences ^5,19^. Mutation of these three residues to alanine (CT-mut1: L337A+L338A+P340A) also abolished CTS2 function without causing other trafficking defects (Fig.6g-h, Fig.S5).

In the immediate juxtamembrane region (aa 326-334), we also introduced the F329A+K330A mutation (CT-mut2), as disruption of this aromatic-basic pair prevents ciliary targeting of Smo and other GPCRs ^24^. Indeed, these residues are also needed for Sstr3’s CTS2 function but not for plasma membrane targeting (Fig.6f-g, Fig.S5).

### Ciliary targeting function of Sstr3-IC3 is mediated by AP[AS]CQ motifs and a basic stretch

After identifying critical residues for CTS2 function of Sstr3-CT, we focused on the CTS1 function of Sstr3-IC3 (aa 231-266). As expected, the CT-mut1 mutation alone did not prevent ciliary targeting when introduced into wild type Sstr3 (Fig.6g,i). We then combined CT-mut1 with seven mutations spanning the entire Sstr3-IC3 (Fig.6d). The first two, IC3-mut2 (VVK231-233AAA) and IC3-Δ1 (Δ234-242), had no effect on Sstr3 CTS1 function (Fig.6g,i).

We then tested the effect of residues 243-255 containing both AP[AS]CQ motifs (243-<u>APSCQ</u>WVQ<u>APACQ</u>-255). IC3-Δ2 (Δ243-247) and IC3-Δ3 (Δ248-255) also had no detectable effect, perhaps due to redundancy between the motifs (Fig.6g,i). To test this, we looked at IC3-Δ4 (Δ243-255). Intriguingly, the IC3-Δ4+CT-mut1 protein was still present in 80% of cilia, which, albeit significantly lower than 100% in CT-mut1, is much higher than 10% in IC3-mut1+CT-mut1, the quadruple phenylalanine mutant (Fig.6g-i). Since both IC3-mut1+CT-mut1 and IC3-Δ4+CT-mut1 readily reach the plasmalemma (Fig.S5), this suggests the bulky phenylalanines in IC3-mut1 have dominant negative effects specifically affecting Sstr3 ciliary targeting.

That is not the whole story, however. Not only did IC3-Δ4 lower the percentage of positive cilia (Fig.6i), but also visibly reduced the amount of ciliary staining (Fig.6j). Quantitation of ciliary signal intensity revealed a 70% decrease of IC3-Δ4+CT-mut1 relative to CT-mut1 control (Fig.6j). Thus, the AP[AS]CQ motifs region does indeed play an important role in Sstr3 CTS1 function.

The second A-Q motif is immediately followed by an arginine-rich stretch (256-RRRRSERR-263), whose deletion (IC3-Δ5) causes a two-fold reduction in both ciliary presence and intensity (Fig.6g-j). Nevertheless, we also noticed that around 40% of IMCD3 cells expressing this mutant fail to traffic it to the cell surface, which may partially or fully explain its reduced ciliary targeting (Fig.6k, Fig.S5). No such intracellular retention was observed with IC3-Δ4+CT-mut1 or CT-mut1 control, where virtually all cells display prominent plasma and/or ciliary membrane localization of these proteins. The last mutant, IC3-mut3, affecting the three residues adjacent to Sstr3’s sixth transmembrane helix, only had a minor effect (Fig.6g-j).

Altogether, these data indicate that CTS1 function in Sstr3-IC3 is encoded by the AP[AS]CQ motif region and the subsequent arginine-rich stretch, even though the latter acts at least partly by enabling transport to the cell surface.

### Ciliary targeting of Htr6 and Sstr3 in neurons also involves CTS1 and CTS2 redundancy

All our ciliary trafficking analyses thus far were performed in the kidney epithelial IMCD3 cell line. However, Htr6 and Sstr3 are above all neuronal GPCRs, with Htr6 being expressed exclusively, and Sstr3 mostly, in the central nervous system, where both receptors accumulate in neuronal cilia ^5,13^. Thus, our IMCD3 data would be of little significance if they were not also applicable to ciliated neurons in the CNS. To test this, we expressed Htr6 and Sstr3, or mutants lacking one or both of our identified CTSs, in primary hippocampal neuron cultures (Fig.7). In these neurons, wild type Htr6 was always clearly detectable in cilia (9 positive cilia out of 9 ciliated and transfected cells: 9/9). For chimera J, lacking CTS1, the proportion was 9/10. For Htr6-(CT-mut3), lacking CTS2, it was 10/10. In contrast, for chimera J-(CT-mut3), lacking both CTS1 and CTS2, the fraction was 2/16 (Fig.7a-b). For Sstr3, the proportions were 9/10 for wild type, 8/10 for IC3-mut1 (lacking CTS1 function), 8/8 for CT-mut1 (lacking CTS2), and 2/12 for the double mutant IC3-mut1+CT-mut1 (Fig.7c-d). Therefore, ciliary targeting of Htr6 and Sstr3 in hippocampal neurons follows the same rules as in IMCD3 cells.

**Figure 7.**
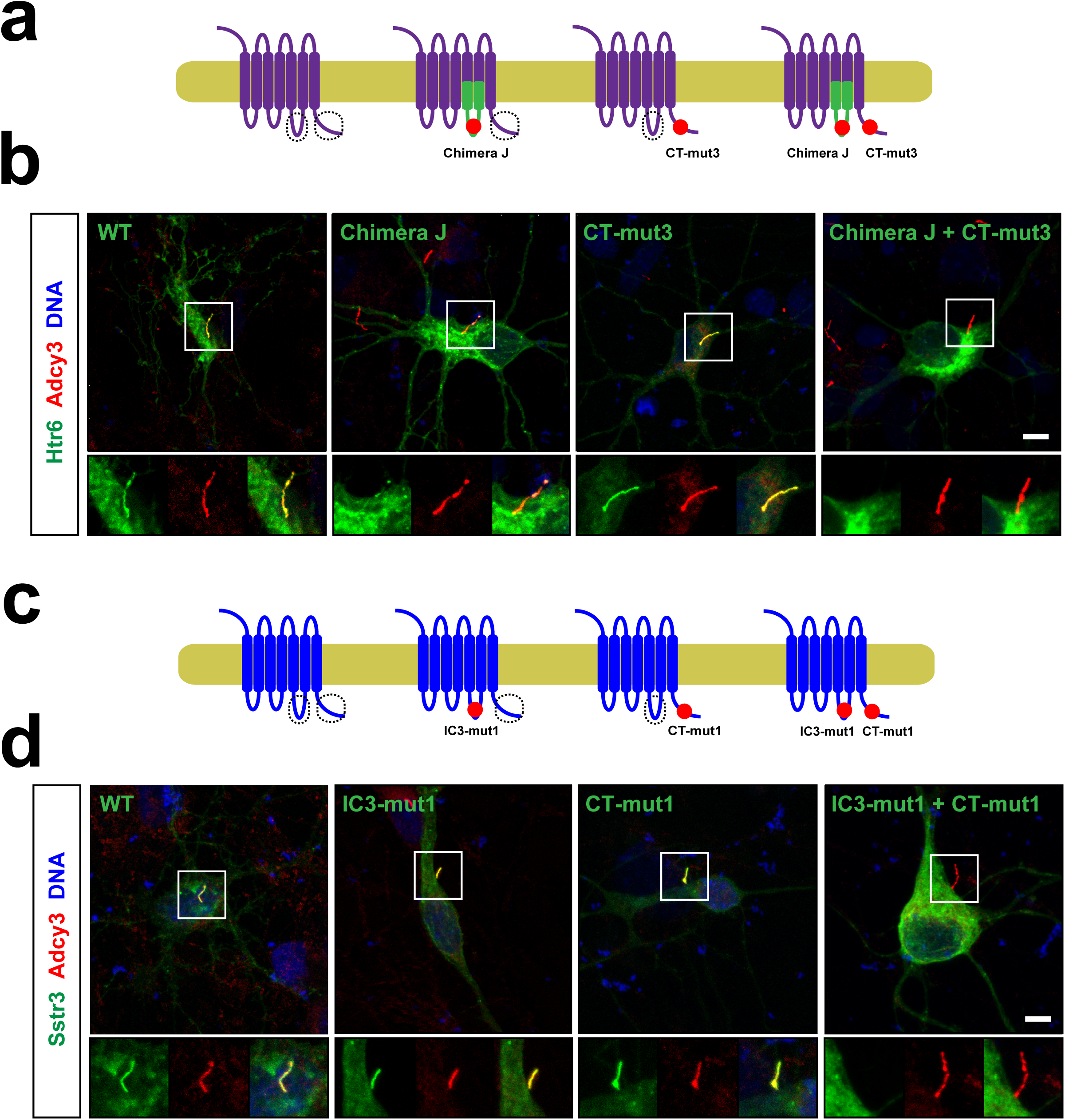
Ciliary targeting of Htr6 and Sstr3 in hippocampal neurons also depends on redundancy between CTS1 and CTS2. **(a)** Schematic of the Htr6 constructs used here. **(b)** Cultured hippocampal neurons expressing C-terminally EGFP-tagged constructs from (a) were analyzed by immunofluorescence with antibodies against adenylate cyclase 3 (Adcy3, red). DNA was stained with DRAQ5 (blue) and EGFP fluorescence was directly visualized. **(c)** Schematic of the Sstr3 constructs used here. **(d)** Cultured hippocampal neurons from (c) were analyzed as in (b). Scale bars, 10 μm.

### Htr6 and Sstr3 CTs associate with Tulp3

Since ciliary accumulation of both Htr6 (Fig.1) and Sstr3 ^25^ depend on Tulp3, which associates to the IC3 loops of two ciliary GPCRs, Gpr161 and Mchr1 ^27^, we then examined whether Htr6 and Sstr3 also interact with Tulp3 through their IC3s and/or CTs. As controls, we used the IC3s and CTs of a non-ciliary GPCR (β2AR) and of the aforementioned Gpr161 ^27^. Since Tulp3 association to ciliary GPCRs has so far only been detected by proximity biotinylation assays, we performed these experiments as previously described, except for the use of an improved biotin ligase, BioID2 (Fig.8a-b) ^27,31^. As expected, Gpr161-IC3 but not β2AR-IC3 robustly associates with Tulp3 (Fig.8c). Interestingly, Htr6-IC3 showed no Tulp3 association, while only a minor one was seen for Sstr3-IC3 (Fig.8c). In contrast, Gpr161-CT showed no specific Tulp3 association when compared to β2AR-CT, whereas both Htr6-CT and Sstr3-CT strongly associated to Tulp3, with Sstr3-CT inducing the strongest Tulp3 biotinylation (Fig.8c). Thus, it appears different ciliary GPCRs associate with Tulp3 in different ways: some rely mostly on their IC3s while others use their CTs.

**Figure 8.**
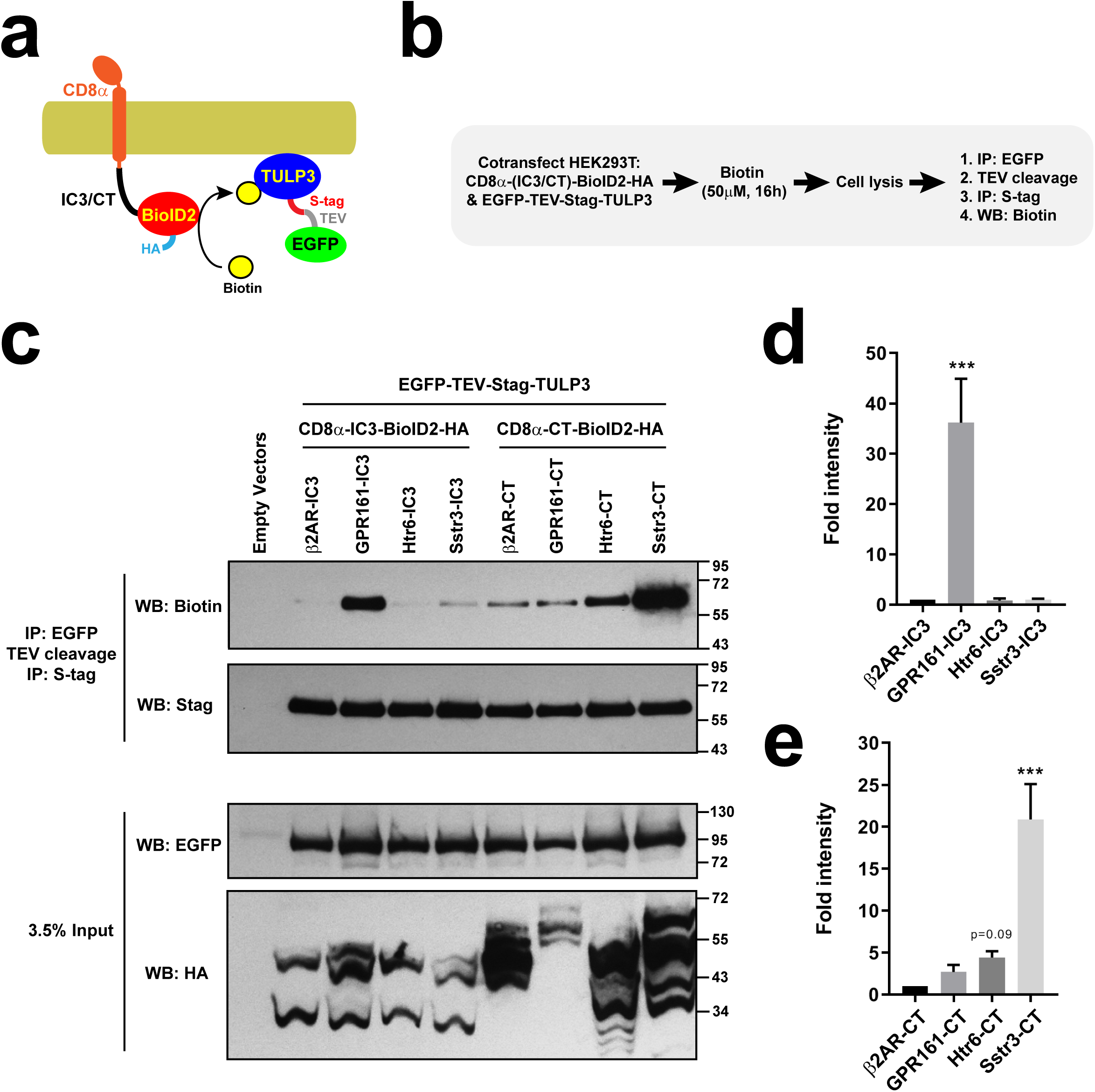
Htr6 and Sstr3 CTs associate with ciliary trafficking adapter Tulp3. **(a-b)** Schematic and protocol of BioID2 proximity labeling assay (PLA). HEK293T cells were cotransfected with plasmids encoding EGFP-TEV-Stag-TULP3 and a fusion protein containing the extracellular and transmembrane regions of CD8α (aa 1-206), the C-terminal tail (CT) or third intracellular loop (IC3) of a GPCR, the BioID2 biotin ligase, and an HA epitope. In presence of biotin (50μM, 16h), BioID2 biotinylates surrounding proteins in a proximity-dependent manner. After cell lysis, TULP3 was affinity purified by two sequential immunoprecipitations (IP) and its biotinylation assessed by Western blot (WB). **(c)** SDS-PAGE and WB analysis of immunoprecipitated S-tagged TULP3 (top two panels) and of the cleared cell lysates (bottom two). In the IPs, Neutravidin-HRP was used to detect TULP3 biotinylation (top) and anti-Stag antibody to detect its total levels. In the lysates, anti-EGFP and anti-HA tag antibodies were used to detect EGFP-TEV-Stag-TULP3 and the CD8 fusions, respectively. Molecular weight markers are indicated on the right (kDa). **(d-e)** Biotinylated TULP3 signal was quantitated from n=4 independent experiments like the one in (c). Biotinylation by IC3 constructs (d) and by CT constructs (e) was normalized relative to β2AR-IC3 and β2AR-CT, respectively, Data are mean ± SEM and were analyzed by one-way ANOVA followed by Dunnett’s multiple comparisons tests. Significance shown as p<0.001 (***).

We also determined which of Tulp3’s two domains is involved in its association with ciliary GPCRs. Tulp3 N-terminal domain, which binds to the IFT-A ciliary trafficking complex ^25^, did not show any association to Htr6-CT (Fig.S6). On the other hand, Tulp3 C-terminal domain, its phosphoinositide-binding Tubby domain, readily associated with Htr6-CT (Fig.S6), consistent with previous data showing that Tulp3 association to ciliary GPCRs requires the former’s ability to bind phosphoinositides ^27^.

### Htr6 CTS2 antagonizes Tulp3 association

We then used proximity biotinylation to test whether CTS2 mutation in Htr6-CT affects Tulp3 association. To our surprise, the CT-mut3 and Δ390-407 mutations, both lacking the LPG motif essential for CTS2 function (Fig.3), displayed much stronger Tulp3 association than wild type Htr6-CT (Fig.9a-b). CT-mut3 also clearly increased Htr6-CT association to Tulp3’s C-terminal Tubby domain (Fig.S6). In contrast, deleting aa 373-389, which are dispensable for cilia localization of chimera J (Fig.3), completely abolished Tulp3 association, as did the bigger Δ373-440 deletion (Fig.9a-b). Thus, aa 373-389 promote Tulp3 binding, whereas aa 390-407 antagonize it. Interestingly, these effects appear to cancel each other out in Htr6-CT(Δ373-407), whose Tulp3 association resembles that of wild type (Fig.9a-b). Finally, Htr6-CT(Δ408-440) also behaves like Htr6-CT(WT), indicating that aa 408-440 are not needed for Tulp3 association with Htr6-CT, even though adding these residues to Htr6-CT(Δ373-440) rescues its Tulp3 association in Htr6-CT(Δ373-407) (Fig.9a-b). Altogether, these data suggest that strong Tulp3 association is not essential for Htr6 ciliary targeting. Instead, preventing excessive Tulp3 association, or promoting its dissociation, may be more important for Htr6 ciliary accumulation.

**Figure 9.**
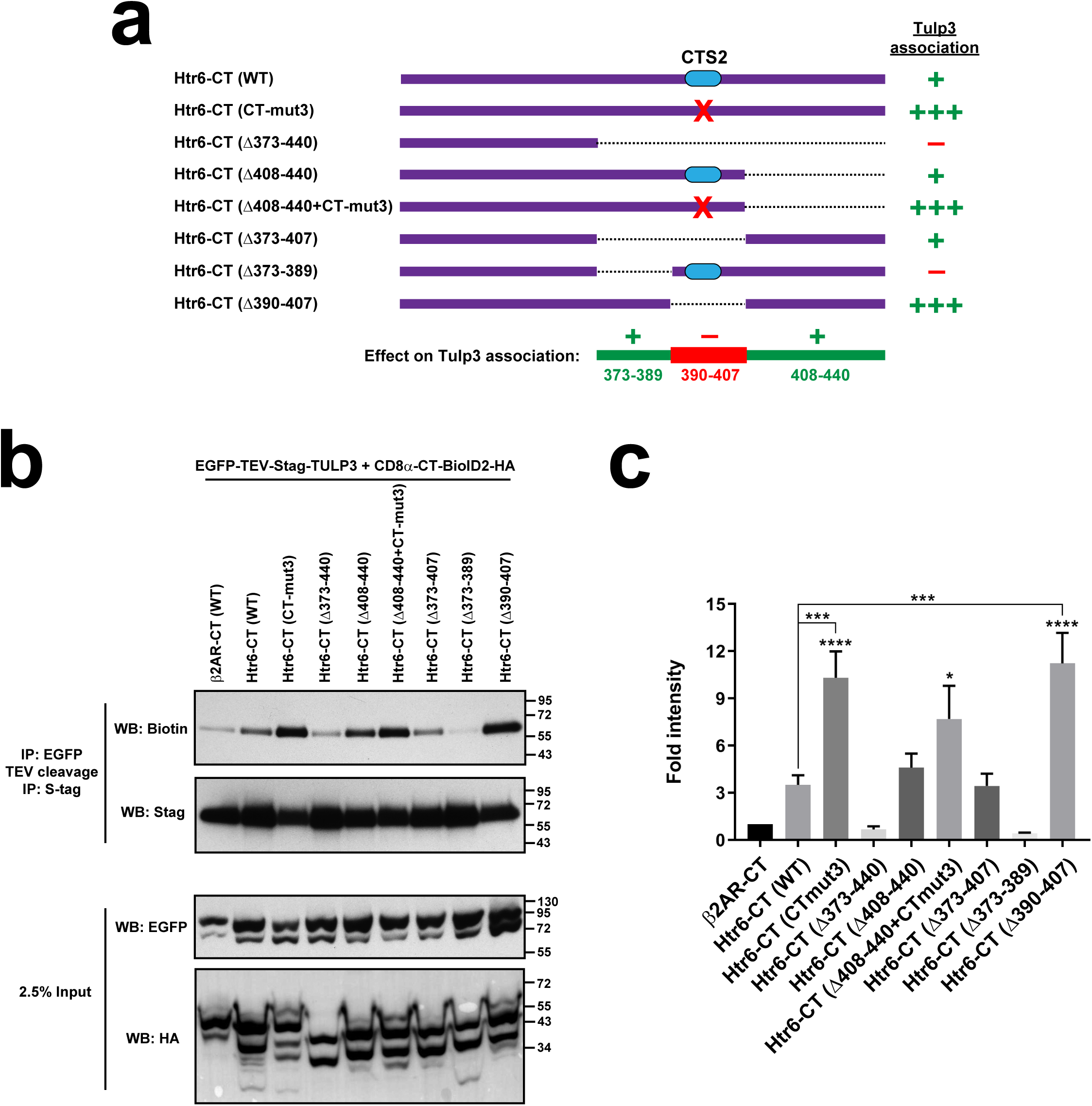
CTS2 of Htr6 antagonizes Tulp3 association to Htr6 C-terminal tail. **(a)** Schematic of the CD8α (aa 1-206)-(Htr6-CT)-BioID2-HA constructs used here, showing only the Htr6-CT moiety. The CTS2 is shown as a blue oval and red crosses indicate the CT-mut3 mutation. Dashed lines indicate deleted regions. The intensity of Tulp3 association is displayed on the right. At bottom, the regions in Htr6-CT promoting Tulp3 association are shown in green, and the CTS2-containing region antagonizing Tulp3 association is shown in red. **(b)** SDS-PAGE and WB analysis of tandem immunoprecipitated S-tagged TULP3 (top two panels) and of cleared cell lysates (bottom two). In the IPs, Neutravidin-HRP was used to detect TULP3 biotinylation (top) and anti-Stag antibody to detect its total levels. In the lysates, anti-EGFP and anti-HA tag antibodies were used to detect EGFP-TEV-Stag-TULP3 and CD8 fusions, respectively. Molecular weight markers on the right (kDa). **(c)** Quantitation of biotinylated TULP3 signal from (b). Data are mean ± SEM (n=7,6,7,5,5,3,5,4,4 independent experiments from left to right, as not all experiments contained all samples). Data were analyzed by one-way ANOVA followed by Tukey’s multiple comparisons tests. Significance shown as p<0.05(*), p<0.001 (***) or p<0.0001(****). Where not explicitly indicated, asterisks represent significance relative to CT-β2AR.

### Htr6 CTS1 and CTS2 contribute to Rabl2 binding

Rabl2 is an atypical Rab small GTPase that promotes anterograde IFT from the ciliary base ^26,32-34^. Recently, we have shown that Rabl2 is required for ciliary targeting of Htr6 and Gpr161, with which it interacts ^26^. Since Htr6 and Rabl2 are readily seen to interact by coimmunoprecipitation, we performed these assays to test whether Htr6 binding to Rabl2 depends on our newly identified CTSs. As expected, EGFP-Rabl2 robustly coimmunoprecipitates wild type myc-Htr6 when both are coexpressed in HEK293T cells (Fig.10a). This was not noticeably affected when myc-Htr6(CT-mut3), lacking CTS2, was used instead. In contrast, coprecipitation of myc-Chimera J, lacking Htr6-IC3 and thus also CTS1, was strongly reduced, and a further reduction was observed with myc-Chimera J (CT-mut3), indicating that CTS2 positively regulates the interaction in absence of Htr6-IC3 (Fig.10a). These data were confirmed by immunoprecipitation of Flag-Rabl2 (Fig.S7).

**Figure 10.**
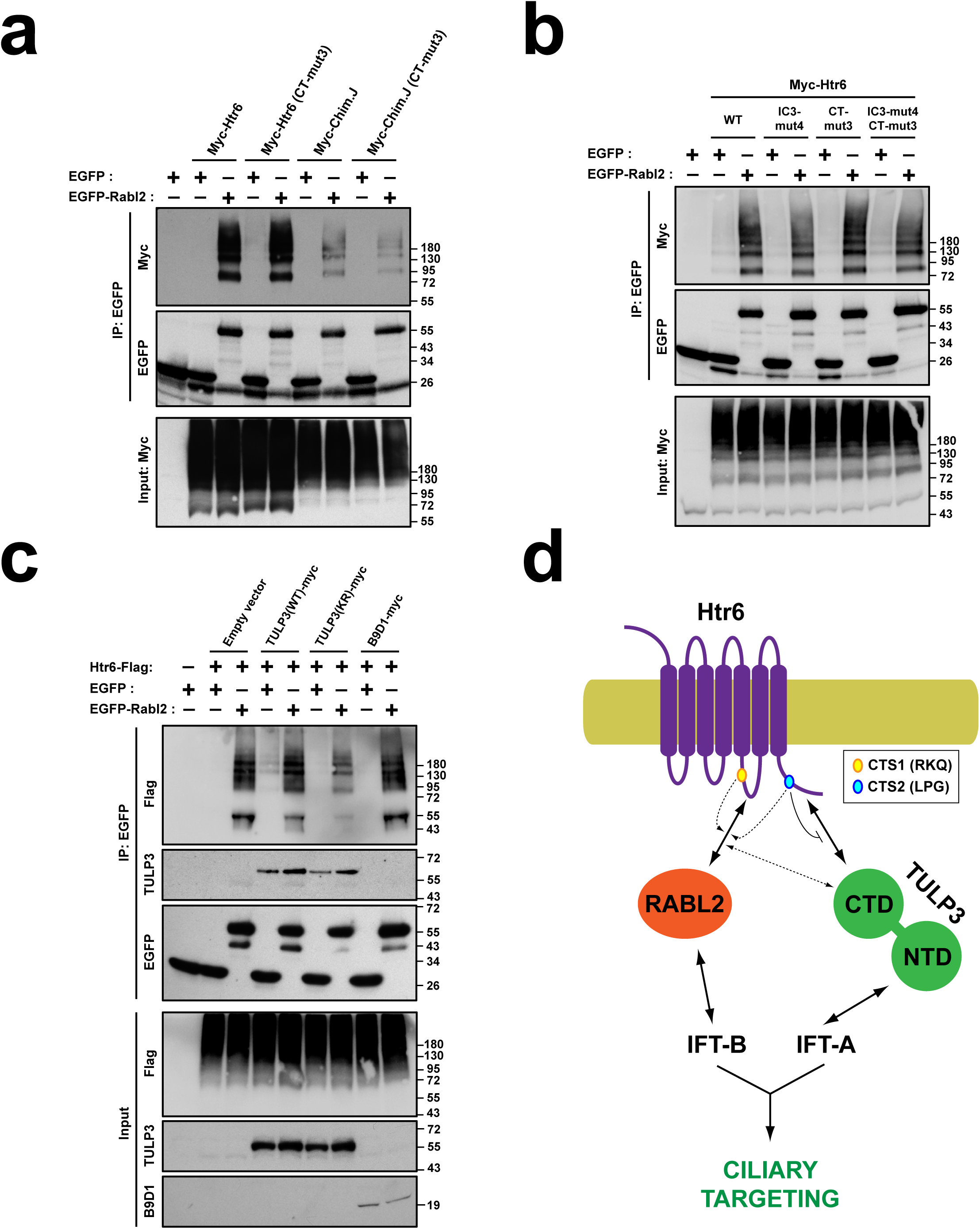
Effects on Htr6-Rabl2 interaction of Htr6-CTSs and TULP3. **(a)** Lysates from HEK293T cells expressing the proteins indicated at the top were immuno-precipitated with anti-EGFP antibodies and analyzed by Western blot with anti-Myc and anti-EGFP antibodies, as indicated. **(b)** Experiment as in (a) except that Htr6-IC3, instead of being entirely replaced by Htr7-IC3 (Chimera J), was mutated in its CTS1 (IC3-mut4: RKQ>AAA). **(c)** Lysates from HEK293T cells expressing the above-indicated proteins were immunoprecipitated with anti-EGFP antibodies and analyzed by Western blot with anti-Flag, anti-TULP3, anti-B9D1 and anti-EGFP antibodies, as indicated. B9D1-myc was used as negative control. TULP3-KR lacks critical lysine and arginine residues needed for phosphoinositide binding. Molecular weight markers are shown on the right of all panels in (a-c). In all cases, input is 2.5% of lysate used for IP. **(d)** Model of Htr6 ciliary targeting. Htr6-CT is the main contributor to TULP3 binding, although CTS2 (with its critical LPG residues) antagonizes that binding, possibly to release Htr6 once inside cilia. TULP3 binding to Htr6 is via its CTD (Tubby domain), as shown in Fig.S6. Htr6-IC3, partly through CTS1 (with its critical RKQ residues), contributes to Rabl2 binding, which CTS2 also promotes in absence of Htr6-IC3 (a). TULP3 may also regulate Rabl2-Htr6 binding. This, however, needs to be established more solidly, like the other effects depicted with dashed lines. Since both RABL2 and TULP3 associate with IFT complexes (see text), these provides a plausible mechanism for Htr6 ciliary targeting.

We then repeated the EGFP-Rabl2 coimmunoprecipitation experiments using myc-Htr6 (IC3-mut4), lacking the critical RKQ motif, instead of myc-Chimera J (Fig.10b). Myc-Htr6 (IC3-mut4) interaction with EGFP-Rabl2 was moderately reduced relative to myc-Htr6 (WT), whose interaction was indistinguishable from myc-Htr6 (CT-mut3), as before. The double mutant, myc-Htr6 (IC3-mut4 + CT-mut3) behaved like the single IC3-mut4 mutant (Fig.10b). These data confirm the positive effect of CTS1 on Rabl2 binding, while suggesting that CTS2’s positive effect is only unveiled in the absence of Htr6-IC3 residues other than those in CTS1.

Finally, we tested whether myc-Tulp3 affects how EGFP-Rabl2 coprecipitates Htr6-Flag (Fig.10c). Coexpression of a negative control protein, myc-B9D1, did not affect Htr6-Rabl2 binding, and neither did coexpression of wild type myc-Tulp3. Reduced Htr6-Flag coprecipitation was instead seen with myc-Tulp3-KR (K268A+R270A), a mutant whose C-terminal Tubby domain cannot bind phosphoinositides (Fig.10c) ^25^. This suggests a possible Tulp3 and phosphoinositide-dependent regulation of Htr6-Rabl2 binding. Of note, both wild type and KR versions of Tulp3 were also coprecipitated by EGFP-Rabl2 (Fig.10c). Altogether, these data suggest several possible connections between Htr6, Rabl2 and Tulp3, which merit further study (Fig.10d).

## DISCUSSION

Herein, we have made several important contributions to our understanding of Htr6 and Sstr3 ciliary targeting. Besides confirming that their IC3s are sufficient but dispensable for ciliary localization ^16,20^, we discovered novel CTSs in the C-terminal tails of these GPCRs. Like the IC3 ones, these novel CTSs suffice for ciliary targeting of transmembrane proteins. As a result, it takes removal of both CTS1 and CTS2 to prevent ciliary targeting of Htr6 or Sstr3.

Initially, the functional redundancy between CTS1 and CTS2 appeared partial, as both chimera D (lacking CTS2) and chimera J (lacking CTS1) localize to cilia less efficiently than wild type Htr6 (Fig.2). Nevertheless, Htr6(CT-mut3), wherein CTS2 is specifically disrupted by point mutations (LLL398-400AAA), localizes to cilia as efficiently as wild type Htr6 (Fig.4), indicating that CTS2 is fully redundant. The fact that CTS1 and CTS2 both drive ciliary accumulation with comparably high efficiency in the CD8α chimera studies suggests redundancy is nearly or fully complete (Fig.5d-f). If so, introducing IC3-mut4 (RKQ216AAA) into wild type Htr6 should not significantly reduce ciliary targeting. Such strong redundancy suggests that loss of ciliary targeting of these receptors is deleterious and evolutionarily selected against. Consistently, mounting evidence indicates that ciliary targeting is critically important for the function of ciliary GPCRs such as Htr6 ^5,6,9,13,14^.

We have also shed light to the role of A-Q motifs. As previously reported ^20^, we have confirmed that chimera N, which contains the first half of Htr6-IC3 in an Htr7 background, localizes to cilia, whereas chimera N (A230F+Q234F), lacking the canonical 230-ATAGQ-234 motif, fails to do so (Fig.S1a-c). However, we have also demonstrated that this AQ>FF mutation strongly impairs chimera N’s plasma membrane targeting, likely explaining why it does not reach cilia (Fig.S1d-e). Furthermore, ciliary targeting of Htr6(CT-mut3) is unperturbed by the same AQ>FF mutation or by the IC3-Δ3 deletion removing the whole A-Q motif (Fig.4a-f). Altogether, this shows that the A-Q motif in Htr6-IC3 is not required for CTS1 function. Accordingly, a canonical Ax[AS]xQ motif is only present in murine Htr6 but not in humans or other mammals, whereas the RKQ motif we identified is conserved across vertebrate Htr6 orthologs (Fig.S7).

Intriguingly, the RKQ motif is preceded by two alanines, making it a non-canonical AxxxQ motif (214-AARKQ-218) ^19,20^. The significance of this is unclear, though, as R216 matters more than Q218 for Htr6 ciliary targeting (Fig.4g-h). On the other hand, alanines 214-215 are fully conserved in vertebrates, but then so are leucines 212-213 and other nearby residues dispensable for CTS1 function (Fig.S7a, Fig.4a,e-f). Checking how the A214F and A215F mutations affect Htr6 intracellular and ciliary trafficking would clarify this issue.

The story is different for Sstr3, which has two A-Q motifs in rodents but only one in humans and other mammals (Fig.S7c). Mutating both A-Q motifs to F-F motifs (IC3-mut1) fully abolishes CTS1 function without affecting plasma membrane trafficking, as seen for instance with the IC3-mut1+CT-mut1 mutant (Fig.6d-h, Fig.S5). However, this is partly due to dominant effects of the added phenylalanines, as IC3-Δ4, which deletes both A-Q motifs (243-<u>APSCQ</u>WVQ<u>APACQ</u>-255), only partially reduces CTS1 function, again without interfering with cell surface expression (Fig.6d-k, Fig.S5). Still, this partial reduction, which affects ciliary intensity more strongly than cilia positivity (Fig.6i-j), indicates that the A-Q motifs in Sstr3 play an important and specific role in ciliary targeting.

Unlike IC3-Δ4, no impairment in CTS1 function is seen with IC3-Δ2 (Δ243-247) and IC3-Δ3 (Δ248-255) (Fig.6i-j), suggesting that both A-Q motifs redundantly promote ciliary targeting of mouse Sstr3. We would expect no such redundancy in human Sstr3, deletion of whose single A-Q motif may suffice to disrupt CTS1 function similar to what IC3-Δ4 does in mouse. This, however, remains untested.

The identity of the tandem AP[AS]CQ motif residues affecting CTS1 function in Sstr3-IC3 also remains unclear. The cysteines are very good candidates, as they are required for Sstr3-IC3 to bind the BBSome, a ciliary cargo adapter required for Sstr3 ciliary accumulation ^30,35^. Furthermore, these cysteines are needed for proper ciliary targeting of CD8α-(Sstr3-IC3) ^30^. Still, the roles of these cysteines have not yet been tested in full length Sstr3. This can now be done by mutating them in combination with CT-mut1 or a similar CTS2-disrupting mutation. The same can be applied to the conserved alanines, prolines and glutamines within the motifs, as they may also have important roles in CTS1 function ^20,23,30^.

In Htr6, we fully mapped the individual residues required for CTS1 and CTS2 function, leading to identification of RKQxxxV and LPG motifs in IC3 and CT, respectively. The former is highly conserved in vertebrates, while the latter is restricted to mammals (Fig.S7a-b). Besides Htr6 orthologs, we also searched these motifs within the sequences of most known ciliary GPCRs (not shown). With a few exceptions (e.g. Gpr88, Gpr161, Gpr63 and EP4 contain LPG motifs in their CTs ^27,36^), the same exact motifs are not present in these other receptors. Still, many of them have similar motifs that could potentially perform the same function (e.g. many have RK or similar motifs in their IC3s). However, in order for these analyses to be more meaningful, we would need to know which residue substitutions preserve or disrupt CTS function at each position within the motif. Thus, these analyses are somewhat premature at this point. As increasing numbers of GPCR CTSs are discovered and characterized, in silico approaches to predict such CTSs should eventually become more reliable.

Our mapping of CTS1 and CTS2 in Sstr3 was less exhaustive than for Htr6, but still very informative. CTS1 function of Sstr3-IC3 relies on the already discussed A-Q motifs, and on an arginine-rich stretch immediately after (256-RRRRSERR-263) (Fig.6i-j). This stretch is very conserved in vertebrates (Fig.S7c), but its effect on ciliary targeting is at least partly due to its role in plasma membrane trafficking, as evidenced by strong intracellular retention of IC3-Δ5+CT-mut1 (Fig.6k, Fig.S5). Higher resolution mutagenic analysis of this stretch may potentially uncouple its effects on cell surface expression from any specific ciliary targeting roles. If such specific roles exist, this would mean that both Htr6-IC3 and Sstr3-IC3 rely on basic residues for CTS1 function, as shown for other ciliary GPCRs like NPY2R ^21^.

In Sstr3-CT, we identified the juxtamembrane region as critical for CTS2 function. Inside this region, an essential role is played by the FK motif (CT-mut2), which is required for BBSome binding and is homologous to Smoothened CTS ^24,37^. Also critical is the LLxP motif disrupted in CT-mut1, which vaguely resembles Htr6’s LPG and Rhodopsin’s VxP ^17,19^. While the FK motif is conserved throughout vertebrates, LLxP is restricted to mammals (Fig.S7d). Nevertheless, the significance of species conservation is unclear, as it is not yet known whether Sstr3 (or Htr6) localizes to non-mammalian cilia. Beyond the juxtamembrane region, deleting Sstr3-CT’s glutamate-rich coiled coil (CT-Δ2) reduces cilia localization by half (Fig.6d-g), prompting the speculative hypothesis that ciliary trafficking of Sstr3 may be reinforced by electrostatic interactions between its E-rich C-terminal coiled coil and its R-rich IC3.

We have also shed some light on the mechanisms of action of these CTSs. Regarding Htr6, we have established that its ciliary targeting is Tulp3-dependent, as shown previously for Sstr3 and other ciliary GPCRs ^22,25,27^. Tulp3 is thought to act as a ciliary trafficking adapter by connecting ciliary membrane cargo to the IFT machinery. Interaction with the latter is dependent on Tulp3’s N-terminal domain, whereas cargo association relies on its C-terminal Tubby domain ^25,27^. Accordingly, we find that Htr6 associates with Tulp3 via its Tubby domain (Fig.S6).

On the GPCR side, we have discovered that Tulp3 association is not always mediated by the IC3 (as previously shown for Mchr1 and Gpr161, and confirmed herein for the latter) but can also be mediated by the CT (as we show for Htr6 and Sstr3 (Fig.8)) ^27^. Furthermore, although Tulp3 binds Htr6-CT, this binding does not require the key CTS2 residues, which antagonize rather than promote this interaction. This suggests that Tulp3 dissociation is a critical step for Htr6 ciliary accumulation. This hypothesis is consistent with other evidence pointing to the following multistep model of Tulp3-mediated ciliary GPCR trafficking: (1) Tulp3 associates with cargo GPCRs at the PI(4,5)P2-rich ciliary base, (2) Tulp3 binding to IFT trains allows it to ferry cargo across the TZ and into the ciliary membrane, (3) low PI(4,5)P2 levels at the ciliary membrane prompt Tulp3-cargo dissociation, and (4) Tulp3 is recycled back across the TZ to the ciliary base, where it can engage in new cycles of cargo transport ^25,27,38-41^. According to this model, failure to release Tulp3 inside the ciliary membrane would prevent Tulp3 recycling and its catalytic action as a ciliary transporter. This could explain the loss of Htr6 ciliary targeting we see with mutations causing excessive Tulp3 binding.

Logically, if Tulp3 dissociation is required for ciliary GPCR targeting, then previous Tulp3 association must also be required. This was already proven for Gpr161, whose IC3 CTS is needed for Tulp3 binding ^27^. For Htr6, we have found that different regions in its CT, both before and after CTS2, promote Tulp3 association, including residues 373-389, whose deletion completely abolishes the interaction (Fig.9). Thus, one would expect that deleting these residues would also prevent Htr6 ciliary accumulation. Although we did not directly check this, we saw that deleting residues 371-378 or 379-391 does not impede Htr6 targeting (Fig.3). However, this may be due to the CTs of these two mutants still binding Tulp3. Alternatively, Htr6 may also bind Tulp3 via sequences other than CT or IC3. Clearly, more work is required to further clarify these issues.

Like Tulp3, Rabl2 functions as an adapter connecting IFT trains to ciliary GPCRs. And like Tulp3, Rabl2 is required for targeting of ciliary GPCRs such as Htr6 and Gpr161 ^26^. Our Rabl2 experiments (Fig.10 and Fig.S7) suggest that Htr6-Rabl2 binding strongly depends on Htr6-IC3, even though other Htr6 regions also appear to be involved. A more systematic mapping of this interaction, including the role of Htr6-CT, is still needed. Within Htr6-IC3, the CTS1 positively affects Rabl2 binding, while the CTS2 in Htr6-CT promotes Htr6-Rabl2 binding only in absence of Htr6-IC3. Thus, CTS1 and CTS2 redundantly promote Htr6-Rabl2 binding, similar to their redundancy in Htr6 ciliary targeting. To what extent redundancy in Rabl2 binding causes redundancy in ciliary targeting is unclear, however. Our data also suggest possible phosphoinositide-dependent effects of Tulp3 on Htr6-Rabl2 binding. These effects merit further investigation, as does the possibility of Rabl2 affecting Htr6-Tulp3 association.

Herein, we have made significant advances in our understanding of how ciliary GPCRs like Htr6 and Sstr3 accumulate in cilia, and how ciliary adapters like Tulp3 and Rabl2 contribute to this process. Moving forward, it will be interesting to see the molecular details of how ciliary trafficking complexes and their adapters interact with GPCRs and their CTSs to mediate ciliary targeting, and to what extent these mechanisms are conserved across diverse GPCRs.

## METHODS

### Reagents and antibodies

Mouse monoclonal antibodies: acetylated Tubulin (Sigma, T7451, IF: 1:10,000), Arl13b (Proteintech, 66739-1-Ig, IF: 1:300; NeuroMab, 75-287, IF: 1:1000) and S-tag (EMD Millipore, MAC112, WB: 1:5000). Rat monoclonal antibody: HA (Chromotek, 7c9, WB: 1:1000). Rabbit polyclonal antibodies: EGFP (Proteintech, 50403-2-AP, IF: 1:200, WB: 1:1000), Flag (Sigma, F7425, WB: 1:1000), adenylyl cyclase III (Santa Cruz, C-20, IF: 1:100), B9D1 (Novus Biologicals, NBP2-84489, WB: 1:1000) and Htr6 ^26^. Goat polyclonal antibodies: γ-Tubulin (Santa Cruz, sc-7396, IF: 1:200). AlexaFluor (AF)-conjugated donkey secondary antibodies from Thermofisher (all used for IF at 1:10,000): AF488 anti-mouse IgG (A21202), AF488 anti-rabbit IgG (A21206), AF555 anti-mouse IgG (A31570), AF594 anti-rabbit IgG (A21207) and AF647 anti-goat IgG (A21447). Also from Thermofisher were HRP-conjugated secondary antibodies (used for WB at 62ng/ml): goat anti-rabbit IgG (A16104), goat anti-mouse IgG (A16072) and donkey anti-rat IgG (A18739). Biotinylated proteins were detected with Neutravidin-HRP (Thermofisher, A2664, 1µg/ml). D-biotin was from Fisher (BP-232-1).

### Plasmids

Details of all plasmids used in this study can be found in Table S1. Htr6-EGFP, Htr7-EGFP, Sstr3-EGFP, Sstr5-EGFP, Htr7[TM5-V241Htr6]-EGFP (chimera N) and the latter’s AQ>FF mutant have been described ^20^, as have EGFP-TEV-Stag-TULP3, EGFP-RABL2B and Flag-RABL2B ^26,27^. Chimeric constructs, internal deletions and missense mutations were generated by overlap extension PCR ^20^. Most other constructs were created by PCR-amplifying the region of interest using primers containing restriction enzyme targets, and mutations where needed. Amplifications were performed with Platinum SuperFi DNA Polymerase (Thermofisher) and resulting PCR products were digested and ligated into desired vectors. Sequences of all plasmids were confirmed by Sanger DNA sequencing (Eurofins Genomics). Primer sequences and PCR conditions are available on request.

### Cell culture and transfection

Murine inner medullary collecting duct 3 (IMCD3) cells, and their HTR6-IMCD3 derivative clone ^26^, were cultured in DMEM/F12 medium supplemented with 10% fetal bovine serum (FBS). Human embryonic kidney 293T (HEK293T) cells were maintained in DMEM with 10% FBS. All cell lines were grown at 37 °C and 5% CO2 in a humidified atmosphere and were mycoplasma-free, as ascertained by regular tests. IMCD3 cells were reverse transfected at ∼50% confluency using JetPrime (Polyplus-transfection) and their cilia analyzed 65 hours later, without serum starvation. RNAi experiments with HTR6-IMCD3 cells are described in next section. HEK293T cells were transfected using either PEI Max (Polysciences) or the calcium phosphate method. Primary hippocampal neurons were cultured as previously described and transfected using Lipofectamine LTX & Plus Reagent methods (Life Technologies/Invitrogen) 7 days after plating ^42^.

### RNA interference

For RNAi, 1×10^5^ HTR6-IMCD3 cells were seeded in 24-well plates, cultured for 24 hours and transfected with 20 pmol siRNA using Lipofectamine RNAiMAX (Thermofisher). Transfected cells were cultured in normal medium for 24h and then serum-starved for 48h before analysis. siRNA oligonucleotides (Sigma) were: siLuc (5’-CGUACGCGGAAUACUUCGAUU-3’), siTulp3#1 (5′-GAAACAAACGUACUUGGAUtt-3′) and siTulp3#2 (5′-GCAGCUAGAAAGCGGAAAAtt-3′).

### Quantitative PCR

Total RNA was isolated from cultured cells using Sepasol (Nacalai Tesque) and reverse transcribed with ReverTra Ace qPCR RT kit (Toyobo). Quantitative PCR was performed using Thunderbird SYBR qPCR mix (Toyobo) and LightCycler96 (Roche). Data were analyzed with ΔΔCt method using siLuc and Gapdh as controls. Primers: Tulp3_F (5’-CAGCTGAAGCTGGACAATCA-3’), Tulp3_R (5’-GGGTTTGGCTGTACCATGAG-3’), Gapdh_F (5’-AACTTTGGCATTGTGGAAGG-3’) and Gapdh_R (5’-TGCAGGGATGATGTTCTGG-3’).

### Immunofluorescence and quantitations

IMCD3 cells were grown on coverslips, fixed 5 min at room temperature (RT) in PBS+4% paraformaldehyde (PFA) followed by 3 min at −20°C in freezer-cold methanol. Cells were then incubated 30–60 min at RT in blocking solution (PBS+0.1% Triton X100+2% donkey serum+0.02% sodium azide), which was also used to dilute primary antibodies. 1 µg/ml DAPI (Thermofisher) was added with secondary antibodies, which were diluted in PBS. After the last round of PBS washes, coverslips were mounted on slides using Prolong Diamond (Thermofisher), incubated overnight at 4°C and imaged with a Leica TCS SP5 confocal microscope. For HTR6-IMCD3, cells were fixed 10 min at RT in PBS+3.7% formalin, permeabilized 10 min in PBS+0.2% Triton X-100 and blocked and stained in PBS+5% BSA. Hoechst33342 (Nacalai Tesque) was used as DNA stain and PermaFluor (Thermofisher) as mounting medium. Cells were imaged using a Zeiss AxioObserver microscope. Primary hippocampal neurons on coverslips were fixed 24h post-transfection in 4% PFA+10% sucrose (15 min, RT), permeabilized in PBS+0.3% Triton X-100+4% donkey serum+1% BSA+0.02% sodium azide (10 min, RT), incubated with anti-Adcy3 antibodies (16-24 h, 4°C), washed thrice in PBS+4% donkey serum+1% BSA+0.02% sodium azide (5 min each), incubated 1h in Alexa Fluor 546-conjugated goat anti-rabbit IgG (1h, RT), washed again and mounted using Immuno-Mount (Thermofisher). Nuclei were visualized with DRAQ5. Samples were imaged on a Leica TCS SP8 laser scanning confocal microscope at the Hunt-Curtis Imaging Facility in the Department of Neuroscience at The Ohio State University. Brightness and contrast of microscopic images were adjusted for optimal visualization with Leica LAS X, Adobe Photoshop or Fiji (Image J). To quantitate percentage of positive cilia for the proteins of interest (GPCRs, chimeras or CD8α fusions), cells that were both transfected and ciliated were counted in seven representative fields spanning the entire coverslip. At least 50 such cells were counted per coverslip and experiment. Cells with no signs of cell surface expression were not counted (such cells were very rare, less than 1%, unless otherwise noted in the text). Quantification of Htr6 intensity in RNAi experiments was performed with Image J as described ^43^. To quantify ciliary signal intensity of Sstr3 mutants, at least 25 cilia per condition were imaged from a representative experiment. Imaging parameters were constant across samples and chosen to avoid signal saturation. Quantitation was then performed on 12-bit images using Fiji to measure average pixel intensity within each cilium and subtracting from it average pixel intensity of an equally sized background region. To quantitate intracellular retention, EGFP-tagged GPCR-transfected cells displaying no plasma membrane or ciliary membrane staining were counted relative to total number of transfected cells.

### Proximity biotinylation assays

Proximity biotinylation experiments were based on ^27^, with some modifications. Briefly, HEK293T cells were cotransfected with plasmids encoding EGFP-TEV-Stag-TULP3 and the proteins of interest fused to BioID2, an improved and smaller biotin ligase ^31^. Transfected cells were grown in presence of 50 µM D-biotin for the last 16 hours before cell lysis. Cells were harvested 48h post-transfection and incubated in lysis buffer (50 mM Tris-HCl, pH 7.4, 200 mM KCl, 1 mM MgCl2, 1 mM EGTA, 10% glycerol, 1 mM DTT, 0.6% IGEPAL CA-630 and Halt protease inhibitor cocktail (Thermofisher, #78429)) at 4°C for 30 min, with rotation. Lysates were then cleared by centrifugation at 4°C for 10 min at 20,000×g. Tulp3 was purified from these lysates by tandem affinity purification. First, EGFP-TEV-Stag-TULP3 was immunoprecipitated with GFP-Trap_MA beads (Chromotek) for 2h at 4°C, with rotation. Immunoprecipitates were then digested for 1h at 25°C with TEV protease (BioVision, #7847) in a buffer containing 50 mM Tris-HCl, pH 8, 100 mM NaCl and 5 mM DTT, according to manufacturer’s instructions. Stag-Tulp3 was then pulled down from the TEV digestion eluates by overnight 4°C rotational incubation using S-protein agarose beads (Merck, #69704). Beads were then eluted with Laemmli buffer and processed for SDS-PAGE in Novex Value 4-20% Tris-Glycine gels (Thermofisher). Proteins were then transferred to nitrocellulose or PVDF membranes and analyzed by immunoblot. For electrochemiluminescent detection, X-ray film or an ImageQuant LAS 500 chemiluminescence CCD camera (GE Life Sciences) were used. Immunoblots were quantified using Fiji/Image J.

### Immunoprecipitation and Western blot

For EGFP-RABL2B immunoprecipitations, cells were harvested 48h post-transfection and lysed in buffer containing 50 mM Tris-HCl pH 7.5, 150 mM NaCl, 0.5 mM EDTA, 1% Igepal CA-630 and Halt protease inhibitor cocktail (Thermofisher, #78429). After rocking 30 min at 4°C in lysis buffer, lysates were cleared by centrifugation at 20,000×g and 4°C for 10 min. For immunoprecipitation with GFP-Trap_MA beads (Chromotek), cleared lysates were rocked for 2h at 4°C. Beads were then eluted with Laemmli buffer and processed for SDS-PAGE in Novex Value 4-20% Tris-Glycine gels (Thermofisher). Proteins were then transferred to nitrocellulose or PVDF membranes and analyzed by immunoblot. For electrochemiluminescent detection, X-ray film or a Fusion Solo S imaging system (Vilber) were used. For Flag-RABL2B immunoprecipitation, cells were lysed at 4°C for 30 minutes in lysis buffer (50 mM Hepes pH 7.5, 150 mM NaCl, 5 mM MgCl2, 0.5% NP-40, 1 mM DTT, 0.5 mM PMSF and 2 µg/ml leupeptin). Lysates were then cleared by centrifugation (17,000 g for 10 min) and their protein content analyzed. Immunoprecipitations were performed on 2 mg of total protein by incubating 2h at 4°C with anti-Flag agarose beads (Sigma). Beads were then washed in lysis buffer and bound polypeptides analyzed by SDS-PAGE and immunoblotting. 20 µg of lysate was loaded in the input lane.

### Statistical analysis

GraphPad Prism 8 software was used to graph and statistically analyze data. Specific details of each experiment are provided in the corresponding figure legends.

## Supporting information

Supplement Barbeito et al 2020

## ACKNOWLEDGMENTS

We thank members of the Garcia-Gonzalo, Kobayashi and Mykytyn labs for useful discussions. This work was supported by European Regional Development Fund (ERDF)-cofunded grants from the Spanish Ministry of Economy and Competitiveness (MINECO) to FRGG (SAF2015-66568-R and RYC2013-14887), by research project grant R21 MH121744 from the NIH/NIMH to KM, and by JSPS KAKENHI (No. 18K06627) to TK. RMM was supported by a MINECO predoctoral grant (BES2016-077828).

## AUTHOR CONTRIBUTIONS

PBG, FRGG, KM and TK designed experiments. PBG, YT, RMM, PM and KM performed experiments under supervision of FRGG, TK and KM. FRGG wrote the manuscript with help from PBG, KM and TK.

## ADDITIONAL INFORMATION

Supplementary Information accompanies this paper (Figures S1-S10, Table S1). Competing interests: The authors declare no competing interests.

