## Supplement Barbeito et al 2020 for "Htr6 and Sstr3 ciliary targeting relies on both IC3 loops and C-terminal tails"

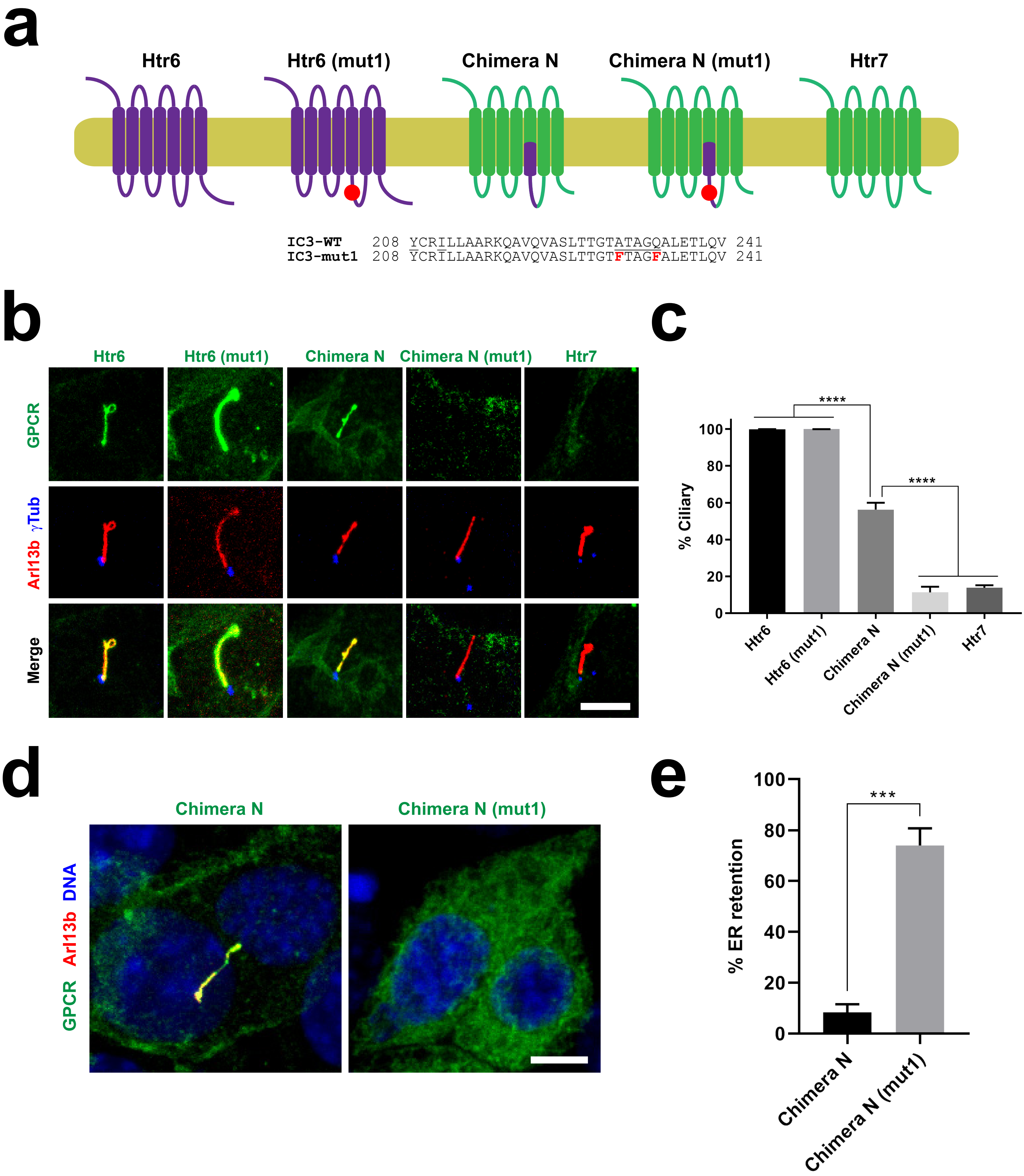


**Figure S1. A-Q motif in Htr6-IC3 is dispensable for ciliary targeting of wild type Htr6. (a)** Schematic representation of Htr7 (green), Htr6 (purple), chimera N, and the mutant versions of the latter two, carrying the A230F+Q234F double mutation (mut1) in the first half of Htr6's IC3 loop, whose sequence is shown below. **(b)** The GPCRs from (a), with EGFP fused to their C-termini, were expressed in IMCD3 cells and their cilia localization was analyzed by immunofluorescence with antibodies against EGFP (green), Arl13b (red) and gamma-tubulin (γTub, blue). Scale bar, 5 μm. **(c)** Percentage of GPCR-positive cilia in GPCR-transfected cells was quantitated from (b). Data are mean ± SEM of n=3 to 5 independent experiments per construct, in each of which at least fifty transfected-cell cilia were counted for each GPCR. Data were analyzed by one-way ANOVA followed by Tukey's multiple comparisons tests. Significance is indicated as $p<0.0001$ (****). **(d)** Immunofluorescence pictures of chimera N and chimera N (mut1) showing the latter's intracellular retention. Scale bar, 5 μm. **(e)** Percentage of transfected cells where indicated chimera was retained intracellularly with no observable plasma membrane staining was quantitated from immunofluorescence experiments. Data are mean ± SEM of n=3 independent experiments, each with at least 150 transfected cells counted per chimera. Significance in unpaired two-tailed Student's t-test shown as $p<0.001$ (***).

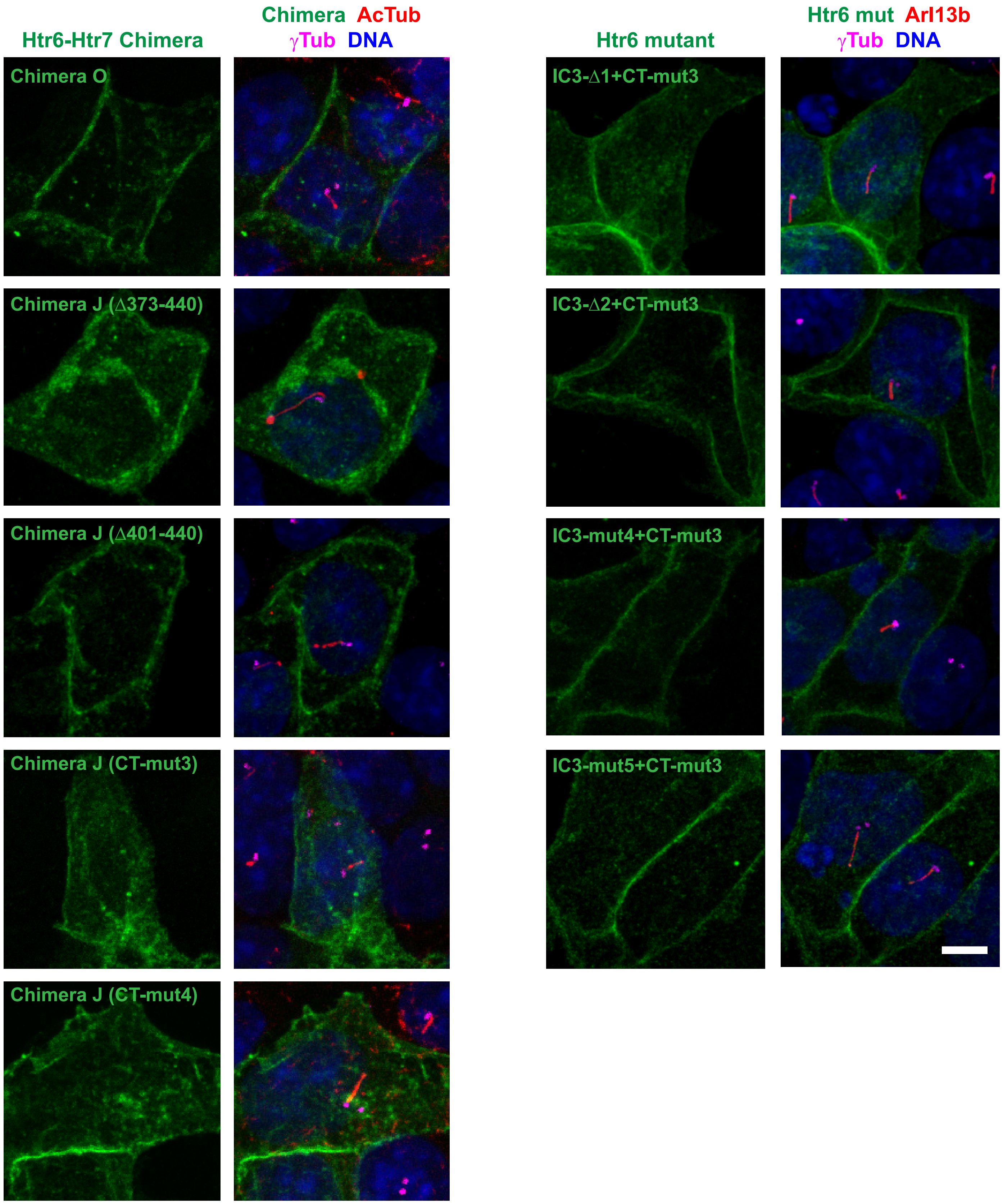


**Figure S2. Plasma membrane targeting of Htr6 mutants. (A)** The indicated Htr6 constructs, all containing C-terminal EGFP, were expressed in IMCD3 cells, which were analyzed by immunofluorescence with antibodies against EGFP (green), acetylated tubulin (AcTub, left) or Arl13b (right) (red) and gamma-tubulin (γTub, magenta). Cells were also stained with DAPI (DNA). All Htr6 constructs clearly label the plasma membrane. Scale bar, 5 μm.

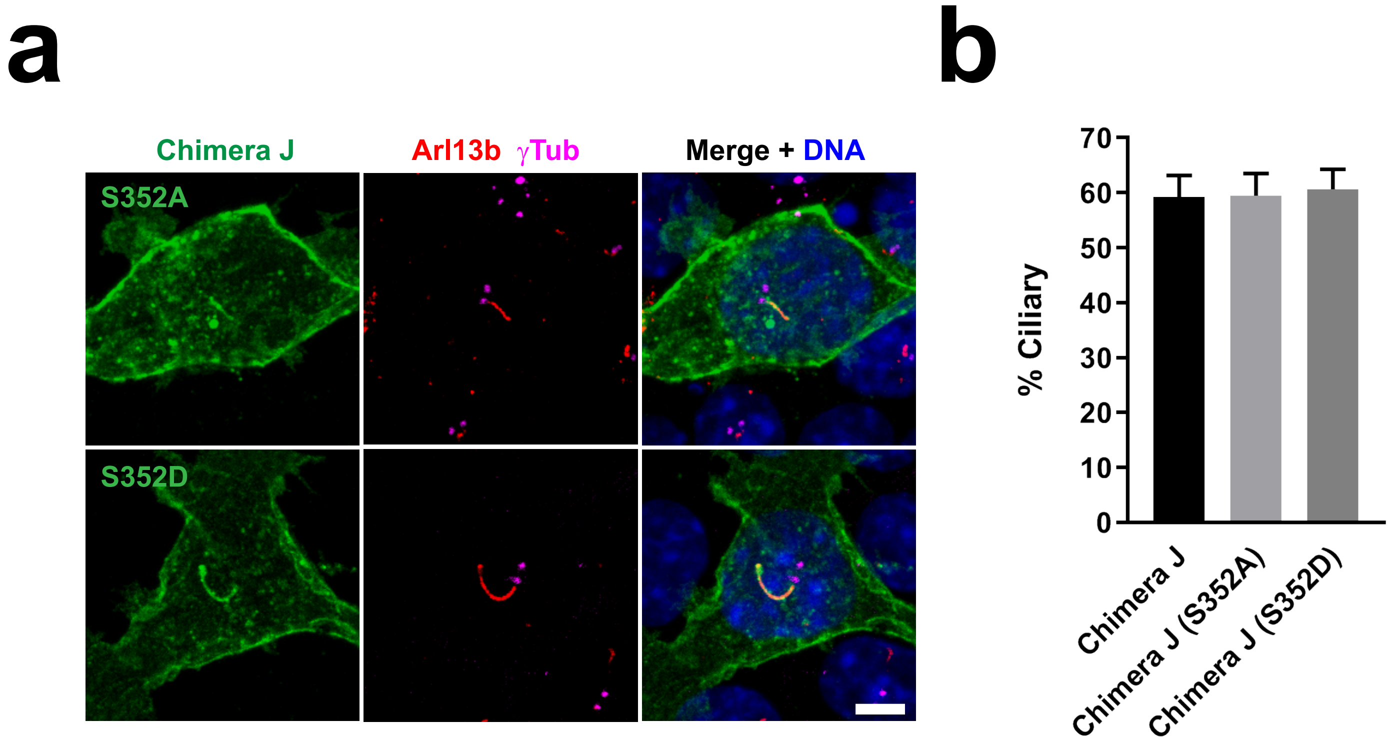


**Figure S3. Cdk5 phosphorylation at Ser-352 in Htr6-CT does not affect Htr6 ciliary targeting. (a)** IMCD3 cells expressing C-terminally EGFP-tagged Chimera J with the S352A or S352D mutations were analyzed by immunofluorescence with antibodies against EGFP (green), Arl13b (red) and gamma-tubulin (γTub, magenta). DNA was stained with DAPI. Arrows indicate cilia. Scale bar, 5 μm. Serine-352 is the mouse Htr6 equivalent of human HTR6 Serine-350, shown to be a target of Cdk5 phosphorylation (Duhr et al. 2014). S352A and S352D are non-phosphorylatable and phosphomimetic Ser-352 mutants, respectively. **(b)** Quantification of ciliary localization from (a). Data are mean ± SEM of n=3 independent experiments per construct, in each of which at least fifty transfected-cell cilia were counted for each GPCR. No significant differences were found by one-way ANOVA.

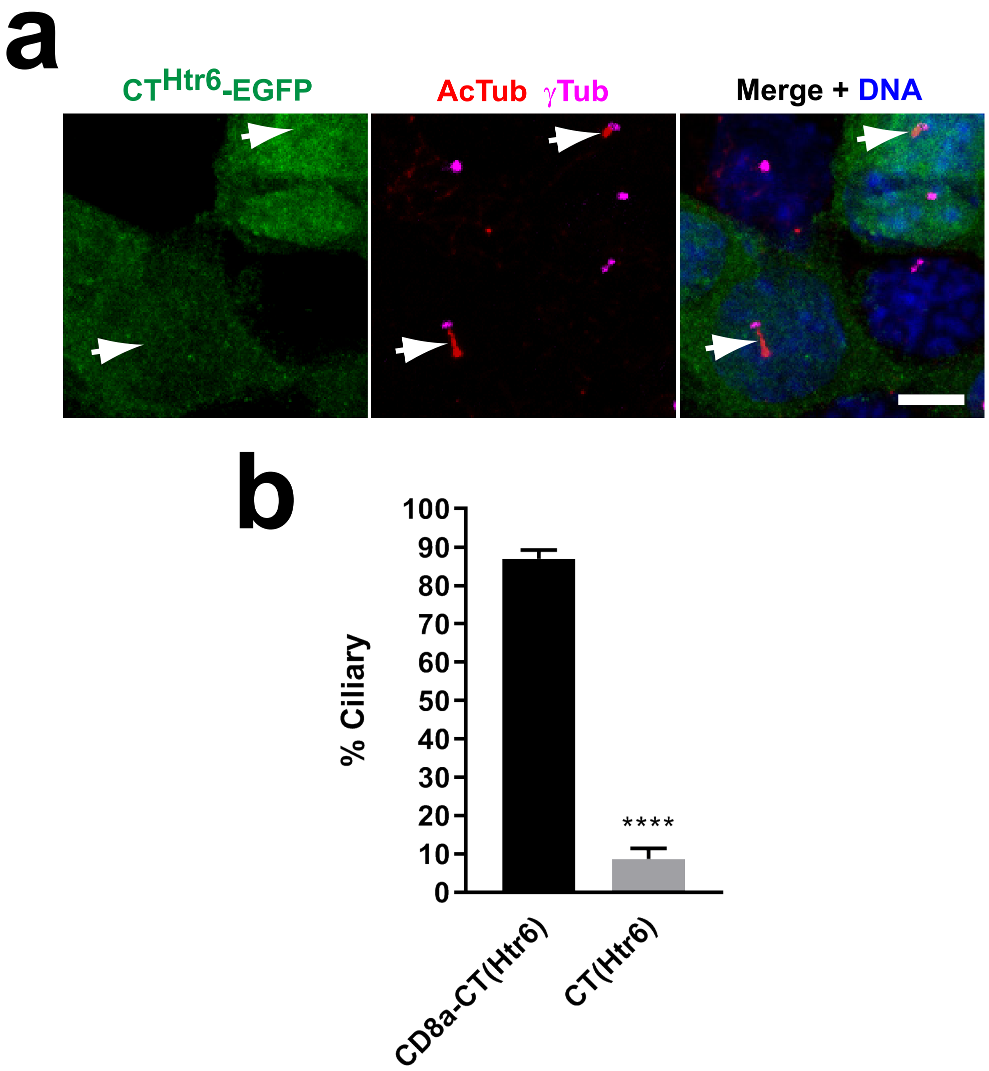


**Figure S4. Membrane association is needed for Htr6-CT to function as a CTS. (a)** IMCD3 cells expressing C-terminally EGFP-tagged Htr6-CT were analyzed by immunofluorescence with antibodies against EGFP (green), acetylated tubulin (AcTub, red) and gamma-tubulin (γTub, magenta). DNA was stained with DAPI. Arrows indicate cilia. Scale bar, 5 μm. (b) Quantification of ciliary localization from (a). Data are mean ± SEM of n=5 and n=3 independent experiments for CD8α(1-206)-CT(Htr6)-EYFP and CT(Htr6)-EGFP, respectively. In each experiment, at least fifty transfected-cell cilia were counted for each construct. Significance in unpaired two-tailed t-test is shown as p<0.0001(****).

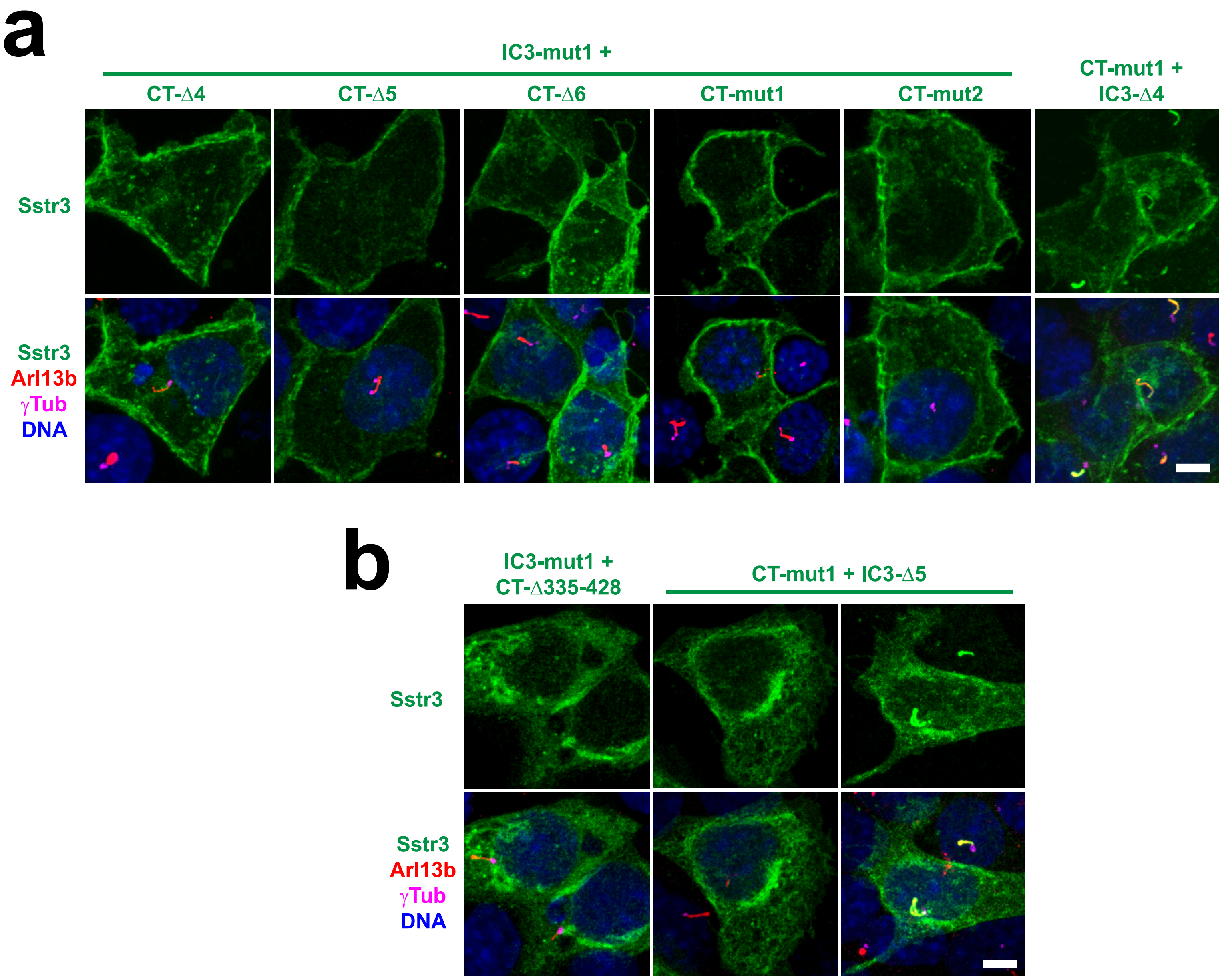


**Figure S5. Plasma membrane targeting of Sstr3 mutants. (a)** The indicated Sstr3 constructs, all containing C-terminal EGFP, were expressed in IMCD3 cells and analyzed by immunofluorescence with antibodies against EGFP (green), Arl13b (red) and gamma-tubulin (γTub, magenta). Cells were also stained with DAPI (DNA). All six constructs clearly reach the plasma membrane. **(b)** Indicated constructs were analyzed as in (a). IC3-mut1+CT-Δ335-428 mutant, lacking all but the first ten residues in Sstr3-CT, consistently accumulates intracellularly and fails to reach plasma and ciliary membranes (left panels). In contrast, CT-mut1+IC3-Δ5, despite its intracellular retention frequency being much higher than normal (middle panels and Fig.6k), can still reach ciliary and/or plasma membrane in about 60% of transfected cells (right panels). Scale bars, 5 μm.

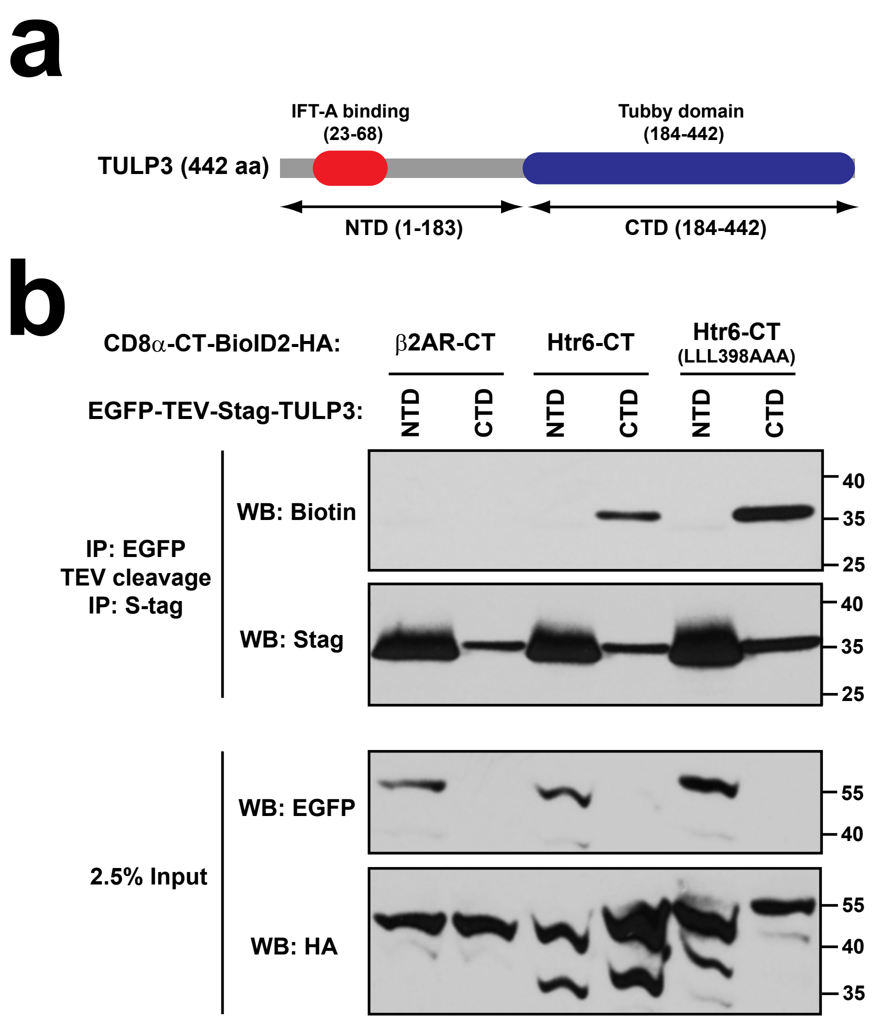


**Figure S6. The C-terminal Tubby domain of Tulp3 is responsible for its association to Htr6-CT. (a)** Schematic of the TULP3 protein depicting its N-terminal (NTD) and C-terminal (CTD) domains. The NTD contains an IFT-A-binding site, whereas the CTD consists of the phosphoinositide-binding Tubby domain. **(b)** Proximity biotinylation assay as done in Figs.8-9. In this case, biotinylation was analyzed for tandem immunoprecipitated S-tagged TULP3-NTD (aa 1-183) or TULP3-CTD (aa 184-442) (top two panels). Analysis of cleared cell lysates is shown in bottom panels. Molecular weight markers are indicated on the right (kDa).

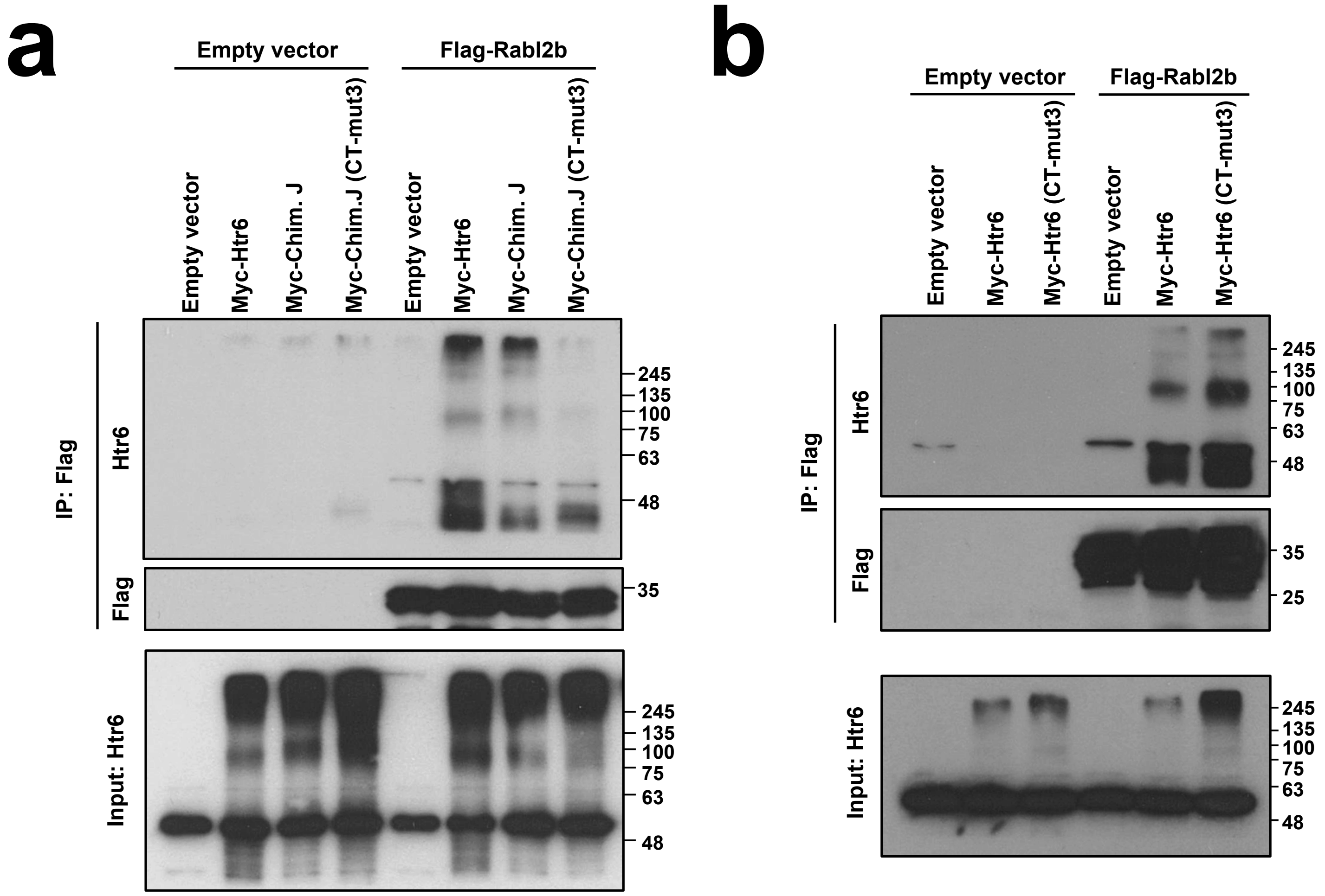


**Figure S7. Mutation of CTS2 reduces Rabl2 binding of Chimera J but not of wild type Htr6**. **(a)** Lysates from HEK293T cells expressing the proteins indicated on top were immunoprecipitated with anti-Flag antibodies and analyzed by Western blot with anti-Htr6 and anti-Flag antibodies, as indicated. **(b)** Experiment as in (a) except that the effect of CT-mut3 was tested in myc-Htr6 rather than myc-ChimeraJ, as indicated. Molecular weight markers are shown on the right of all panels.

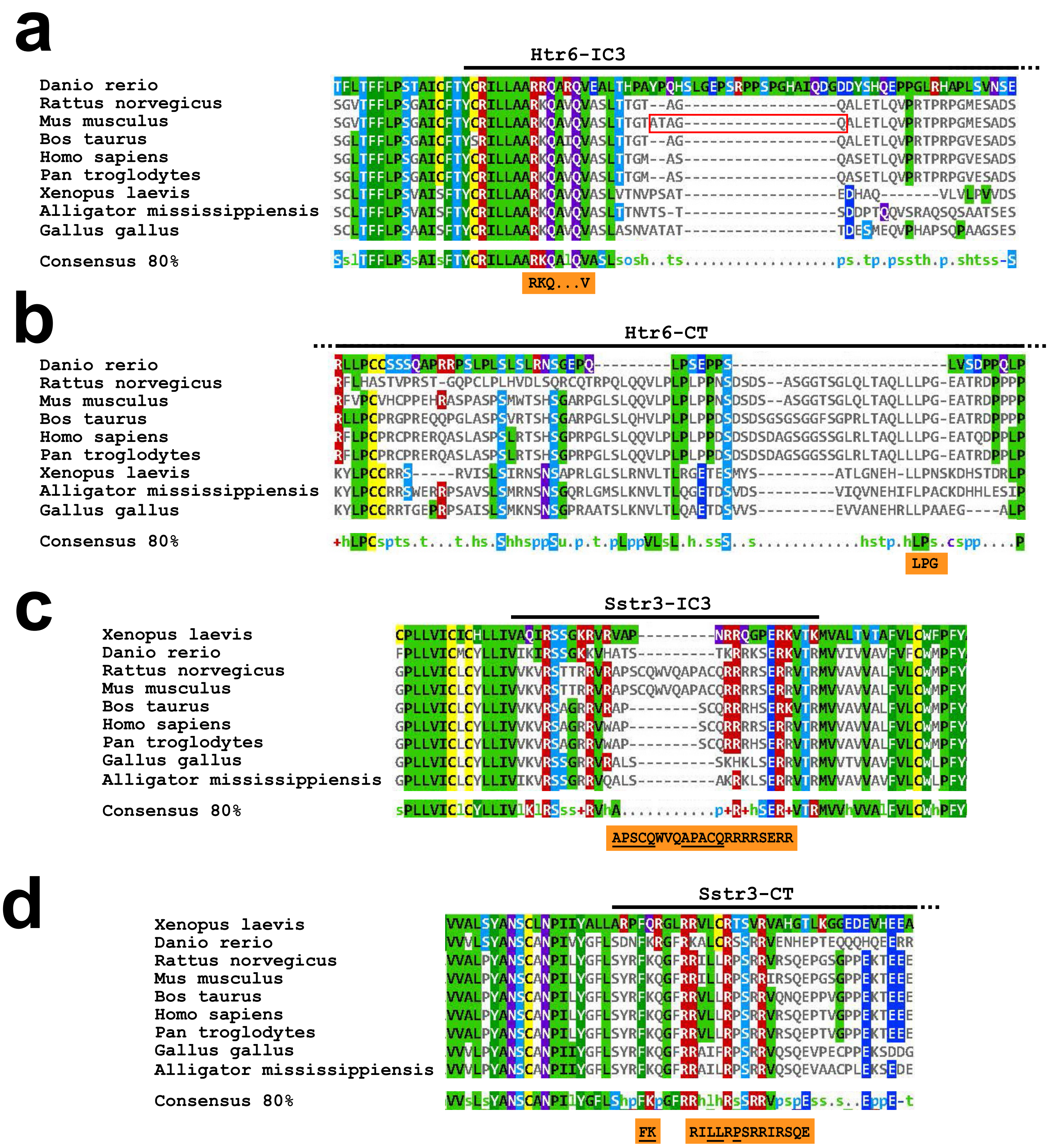


**Figure S8. Evolutionary conservation of Htr6 and Sstr3 ciliary targeting sequences**. **(a)** Htr6 orthologs from indicated species were aligned using Clustal Omega and the resulting alignment visualized with MView. The displayed region includes the first half of Htr6-IC3, as indicated on top. The RKQ...V motif we identified in mouse Htr6 is shown in the orange box below and the mouse-specific ATAGQ motif is marked with a red rectangle. The 80% consensus sequence is also shown. **(b)** Htr6 orthologs were aligned and visualized as in (a) and the C-terminal tail region containing the LPG motif (orange box) is shown here. **(c)** Sstr3 orthologs were aligned and visualized as in (a-b) and the C-terminal tail region containing the LPG motif (orange box) is shown here. The orange box shows the residues we found involved in ciliary targeting of mouse Sstr3, with A-Q motifs underlined. **(d)** Sstr3 orthologs were aligned and visualized as in (c). The juxtamembrane region of Sstr3-CT is shown. The regions we found to affect mouse Sstr3 ciliary targeting are shown on the orange box below, with the FK and LLxP motifs underlined. Sequence identifiers for Htr6 orthologs are: tr|E9QFU3|, tr|A0A0G2JYX7|, sp|Q9R1C8|, tr|E1BE22|, sp|P50406|, sp|Q5IS65|, tr|A0A1L8FPD2|, tr|A0A151MJI5| and tr|D2XUT1| for zebrafish, rat, mouse, bovine, human, chimp, toad, alligator and chick, respectively. For Sstr3 orthologs they are: tr|A0A1L8GH74|, tr|A2BHH9|, sp|P30936|, sp|P30935|, tr|G3MWT4|, sp|P32745|, tr|A0A2J8QPP4|, tr|Q4ZJF2| and tr|A0A151NHS7| for toad, zebrafish, rat, mouse, bovine, human, chimp, chick and alligator, respectively. The lowercase letters in the consensus rows stand for: hydrophobic (h), small (s), aliphatic (l), polar (p), turnlike (t), tiny (u), charged (c), alcohol (o) and positive (+).

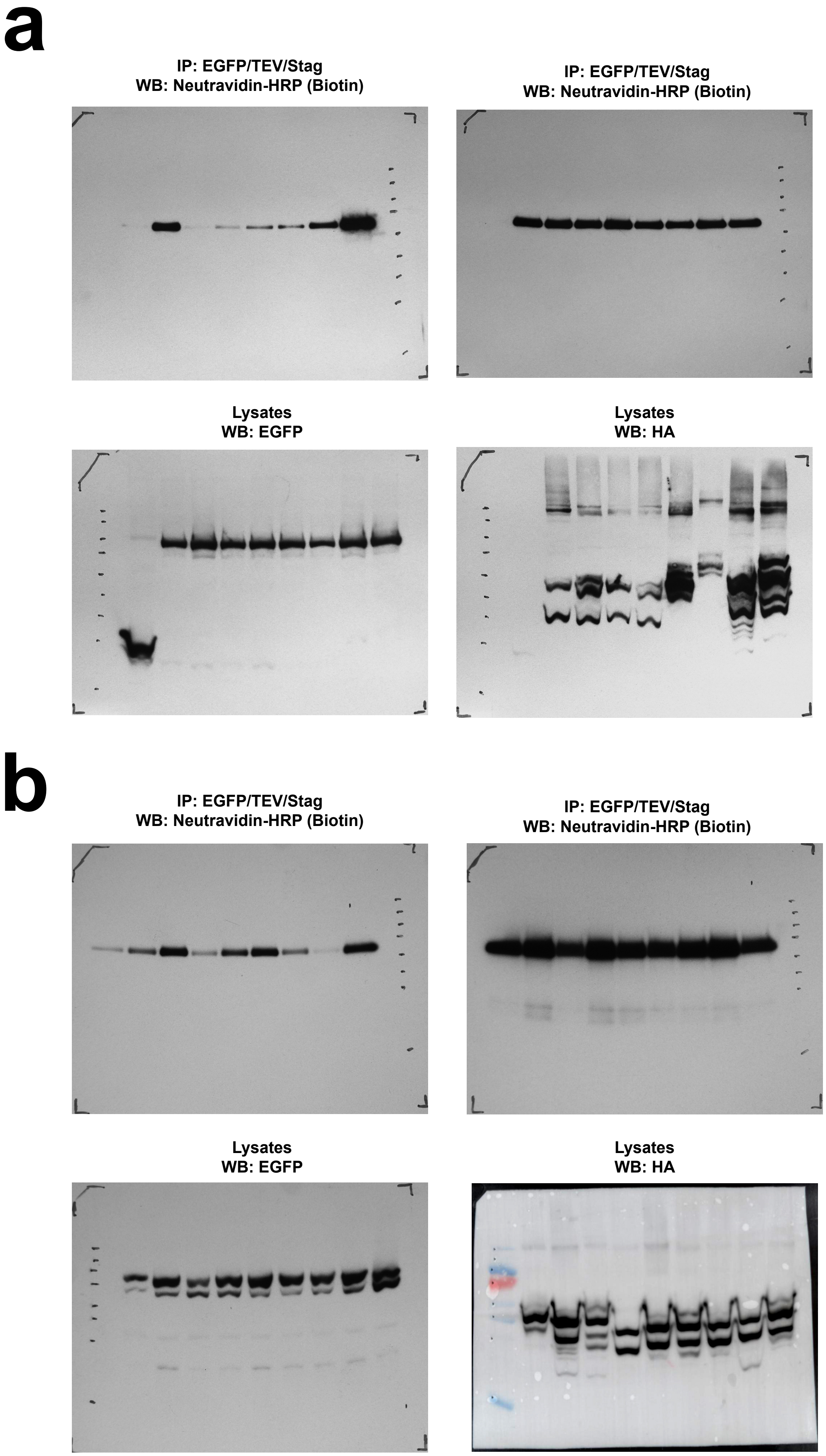


**Figure S9. Full Western blot panels from Figures 8-9**. **(a)** Full panels from Figure 8. **(b)** Full panels from Figure 9.

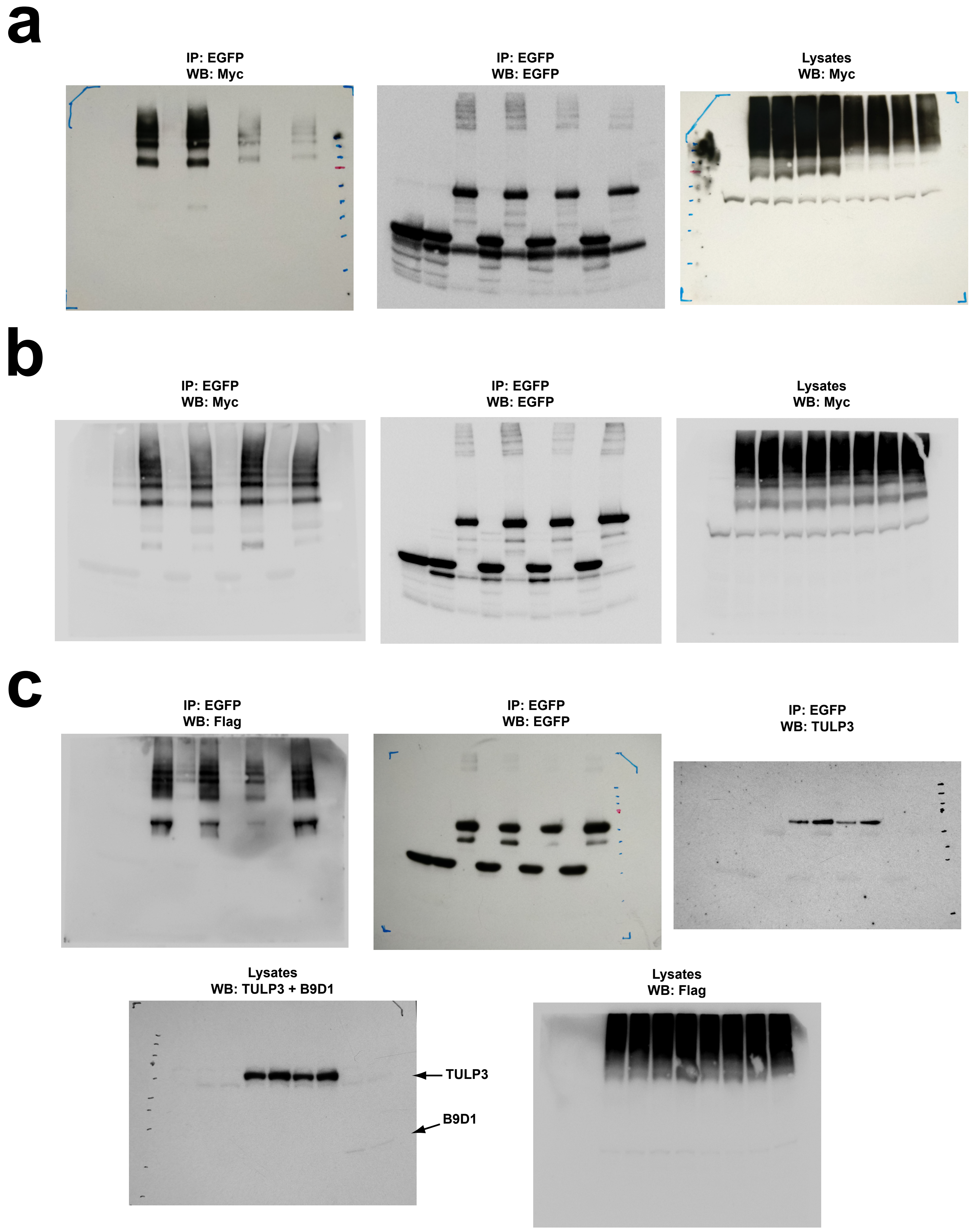


**Figure S10. Full Western blot panels from Figure 10. (a)** Full panels from Figure 10a. **(b)** Full panels from Figure 10b. **(c)** Full panels from Figure 10c.

### Supplementary Table I. Plasmids

| PLASMID NAME | BACKBONE | INSERT | CLONING | SOURCE |
|---|---|---|---|---|
| Htr6-EGFP | pEGFP-N1 | Mouse Htr6 (NP_067333.1) (440 aa) | XhoI-HindIII | This paper |
| Htr7-EGFP | pEGFP-N3 | Mouse Htr7 (NP_032341.2) (448 aa) | EcoRI-BamHI | Berbari et al. 2008a |
| Chimera D-EGFP | pEGFP-N3 | Htr6(1-282)-Htr7(343-448) | EcoRI-BamHI | This paper |
| Chimera J-EGFP | pEGFP-N3 | Htr6(1-198)-Htr7(253-342)-Htr6(283-440) | EcoRI-BamHI | This paper |
| Chimera N-EGFP | pEGFP-N3 | Htr7(1-252)-Htr6(199-241)-Htr7(296-448) | EcoRI-BamHI | Berbari et al. 2008a |
| Chimera O-EGFP | pEGFP-N3 | Htr6(1-198)-Htr7(253-342)-Htr6(283-308)-Htr7(374-448) | EcoRI-BamHI | This paper |
| Chimera Q-EGFP | pEGFP-N3 | Htr6(1-198)-Htr7(253-342)-Htr6(283-308)-Htr7(374-435)-Htr6(369-440) | EcoRI-BamHI | This paper |
| Chimera R-EGFP (a.k.a. Htr7-CT(Htr6)-EGFP) | pEGFP-N3 | Htr7(1-448)-[GS]-Htr6(326-440) (GS linker is BamHI site) | EcoRI-BamHI | This paper |
| Chimera J (Δ369-370)-EGFP | pEGFP-N3 | Chimera J + (Δ369-370) | EcoRI-BamHI | This paper |
| Chimera J (Δ371-378)-EGFP | pEGFP-N3 | Chimera J + (Δ371-378) | EcoRI-BamHI | This paper |
| Chimera J (Δ379-391)-EGFP | pEGFP-N3 | Chimera J + (Δ379-391) | EcoRI-BamHI | This paper |
| Chimera J (Δ373-440)-EGFP | pEGFP-N3 | Chimera J + (Δ373-440) | EcoRI-BamHI | This paper |
| Chimera J (Δ401-440)-EGFP | pEGFP-N3 | Chimera J + (Δ401-440) | EcoRI-BamHI | This paper |
| Chimera J (Δ425-440)-EGFP | pEGFP-N3 | Chimera J + (Δ425-440) | EcoRI-BamHI | This paper |
| Chimera J (mut1)-EGFP | pEGFP-N3 | Chimera J + (LQL392-394AAA) | EcoRI-BamHI | This paper |
| Chimera J (mut2)-EGFP | pEGFP-N3 | Chimera J + (TAQ395-397AAA) | EcoRI-BamHI | This paper |
| Chimera J (mut3)-EGFP | pEGFP-N3 | Chimera J + (LLL398-400AAA) | EcoRI-BamHI | This paper |
| Chimera J (mut4)-EGFP | pEGFP-N3 | Chimera J + (PGE401-403AAA) | EcoRI-BamHI | This paper |
| Chimera J (mut5)-EGFP | pEGFP-N3 | Chimera J + (TRD405-407AAA) | EcoRI-BamHI | This paper |
| Chimera J (mut6)-EGFP | pEGFP-N3 | Chimera J + (PPPP408-411AAAA) | EcoRI-BamHI | This paper |
| Chimera J (mut7)-EGFP | pEGFP-N3 | Chimera J + (TRAPT412-416AAAAA) | EcoRI-BamHI | This paper |
| Chimera J (mut8)-EGFP | pEGFP-N3 | Chimera J + (VVNF417-420AAAA) | EcoRI-BamHI | This paper |
| Chimera J (mut9)-EGFP | pEGFP-N3 | Chimera J + (FVTD421-424AAAA) | EcoRI-BamHI | This paper |
| Chimera J (L398A)-EGFP | pEGFP-N3 | Chimera J + (L398A) | EcoRI-BamHI | This paper |
| Chimera J (L399A)-EGFP | pEGFP-N3 | Chimera J + (L399A) | EcoRI-BamHI | This paper |
| Chimera J (L400A)-EGFP | pEGFP-N3 | Chimera J + (L400A) | EcoRI-BamHI | This paper |
| Chimera J (P401A)-EGFP | pEGFP-N3 | Chimera J + (P401A) | EcoRI-BamHI | This paper |
| Chimera J (G402A)-EGFP | pEGFP-N3 | Chimera J + (G402A) | EcoRI-BamHI | This paper |
| Chimera J (E403A)-EGFP | pEGFP-N3 | Chimera J + (E403A) | EcoRI-BamHI | This paper |
| Chimera N (AQ>FF)-EGFP | pEGFP-N3 | Chimera N + (A230F+Q234F) | EcoRI-BamHI | This paper |
| Htr6 (CTmut3)-EGFP | pEGFP-N1 | Htr6 + (LLL398AAA) | XhoI-HindIII | This paper |
| Htr6 (IC3Δ1+CTmut3)-EGFP | pEGFP-N1 | Htr6 (CTmut3) + (Δ208-219) | XhoI-HindIII | This paper |
| Htr6 (IC3Δ2+CTmut3)-EGFP | pEGFP-N1 | Htr6 (CTmut3) + (Δ220-229) | XhoI-HindIII | This paper |
| Htr6 (IC3Δ3+CTmut3)-EGFP | pEGFP-N1 | Htr6 (CTmut3) + (Δ230-241) | XhoI-HindIII | This paper |
| Htr6 (IC3mut1)-EGFP | pEGFP-N1 | Htr6 (A230F+Q234F) | XhoI-HindIII | This paper |
| Htr6 (IC3mut1+CTmut3)-EGFP | pEGFP-N1 | Htr6 (CTmut3) + (A230F+Q234F) | XhoI-HindIII | This paper |
| Htr6 (IC3mut2+CTmut3)-EGFP | pEGFP-N1 | Htr6 (CTmut3) + (CR209-210AA) | XhoI-HindIII | This paper |
| Htr6 (IC3mut3+CTmut3)-EGFP | pEGFP-N1 | Htr6 (CTmut3) + (LL212-213AA) | XhoI-HindIII | This paper |
| Htr6 (IC3mut4+CTmut3)-EGFP | pEGFP-N1 | Htr6 (CTmut3) + (RKQ216-218AAA) | XhoI-HindIII | This paper |
| Htr6 (IC3mut5+CTmut3)-EGFP | pEGFP-N1 | Htr6 (CTmut3) + (VQV220-222AAA) | XhoI-HindIII | This paper |
| Htr6 (IC3mut6+CTmut3)-EGFP | pEGFP-N1 | Htr6 (CTmut3) + (SLT224-226AAA) | XhoI-HindIII | This paper |
| Htr6 (IC3mut7+CTmut3)-EGFP | pEGFP-N1 | Htr6 (CTmut3) + (TGT227-229AAA) | XhoI-HindIII | This paper |
| Htr6 (IC3mut8+CTmut3)-EGFP | pEGFP-N1 | Htr6 (CTmut3) + (R216A) | XhoI-HindIII | This paper |
| Htr6 (IC3mut9+CTmut3)-EGFP | pEGFP-N1 | Htr6 (CTmut3) + (K217A) | XhoI-HindIII | This paper |
| Htr6 (IC3mut10+CTmut3)-EGFP | pEGFP-N1 | Htr6 (CTmut3) + (Q218A) | XhoI-HindIII | This paper |
| Htr6 (IC3mut11+CTmut3)-EGFP | pEGFP-N1 | Htr6 (CTmut3) + (A219F) | XhoI-HindIII | This paper |
| Htr6 (IC3mut12+CTmut3)-EGFP | pEGFP-N1 | Htr6 (CTmut3) + (V220A) | XhoI-HindIII | This paper |
| Htr6 (IC3mut13+CTmut3)-EGFP | pEGFP-N1 | Htr6 (CTmut3) + (Q221A) | XhoI-HindIII | This paper |
| Htr6 (IC3mut14+CTmut3)-EGFP | pEGFP-N1 | Htr6 (CTmut3) + (V222A) | XhoI-HindIII | This paper |
| Htr6 (IC3mut15+CTmut3)-EGFP | pEGFP-N1 | Htr6 (CTmut3) + (RK216-217AA) | XhoI-HindIII | This paper |
| Htr6 (IC3mut16+CTmut3)-EGFP | pEGFP-N1 | Htr6 (CTmut3) + (R216A+Q218A) | XhoI-HindIII | This paper |
| CT(Htr6)-EGFP | pEGFP-N3 | Htr6 (326-440) | XhoI-BamHI | This paper |
| Myc-Htr6 | pEGFP-N1 | Myc-Htr6 (EGFP removed) | XhoI-NotI | This paper |
| Myc-Htr6 (IC3mut4) | pEGFP-N1 | Myc-Htr6 + (RKQ216AAA) | XhoI-NotI | This paper |
| Myc-Htr6 (CTmut3) | pEGFP-N1 | Myc-Htr6 + (LLL398AAA) | XhoI-NotI | This paper |
| Myc-Htr6 (IC3mut4+CTmut3) | pEGFP-N1 | Myc-Htr6 + (RKQ216AAA+LLL398AAA) | XhoI-NotI | This paper |
| Myc-Chimera J | pEGFP-N1 | Myc-Chimera J (replacing EGFP) | XhoI-NotI | This paper |
| Myc-Chimera J (CTmut3) | pEGFP-N1 | Myc-Chimera J + (LLL398AAA) | XhoI-NotI | This paper |
| CD8a (1-206)-EYFP | pEYFP-N1 | CD8α (1-206) (from CD8A-EGFP (Addgene #86051)) | XhoI-EcoRI | This paper |
| CD8a-(Htr6-IC3)-EYFP | CD8a (1-206)-EYFP | [SAGG]-Htr6 (208-268)-[SAGG] | EcoRI-BamHI | This paper |
| CD8a-(Htr6-CT)-EYFP | CD8a (1-206)-EYFP | Htr6 (326-440) | EcoRI-BamHI | This paper |
| CD8a-(Htr7-CT)-EYFP | CD8a (1-206)-EYFP | Htr7 (390-448) | EcoRI-BamHI | This paper |
| CD8a-(Sstr3-CT)-EYFP | CD8a (1-206)-EYFP | Mouse Sstr3 (NP_033244.2) (aa 325-428) | EcoRI-AgeI | This paper |
| CD8a-(Sstr3-IC3)-EYFP | CD8a (1-206)-EYFP | [SAGG]-Sstr3 (231-266)-[SAGG] | EcoRI-BamHI | This paper |
| CD8a(1-206)-BioID2-HA | MCS-BioID2-HA | CD8α (1-206) + XhoI site | EcoRI-BamHI | This paper |
| CD8a-(Htr6-CT)-BioID2-HA | CD8a(1-206)-BioID2-HA | Htr6 (326-440) | XhoI-BamHI | This paper |
| CD8a-(Htr6-IC3)-BioID2-HA | CD8a(1-206)-BioID2-HA | Htr6 (208-268) (replacing CT(Htr6)) | XhoI-BamHI | This paper |
| CD8a-(B2AR-CT)-BioID2-HA | CD8a(1-206)-BioID2-HA | Human B2AR (NP_000015.1) (aa 330-413) | XhoI-BamHI | This paper |
| CD8a-(B2AR-IC3)-BioID2-HA | CD8a(1-206)-BioID2-HA | B2AR (aa 224-274) | XhoI-BamHI | This paper |
| CD8a-(GPR161-CT)-BioID2-HA | CD8a(1-206)-BioID2-HA | Human GPR161 (NP_722561.1) (aa 328-529) | XhoI-BamHI | This paper |
| CD8a-(GPR161-IC3)-BioID2-HA | CD8a(1-206)-BioID2-HA | GPR161 (aa 213-267) | XhoI-BamHI | This paper |
| CD8a-(Sstr3-CT)-BioID2-HA | CD8a-CT(Sstr3)-EYFP | BioID2-HA (replacing EYFP) | AgeI-NotI | This paper |
| CD8a-(Sstr3-IC3)-BioID2-HA | CD8a(1-206)-BioID2-HA | Sstr3 (231-266) | XhoI-BamHI | This paper |
| CD8a-(Htr6-CT)-BioID2-HA (Δ373-440) | CD8a(1-206)-BioID2-HA | Htr6 (326-440) + (Δ373-440) | XhoI-BamHI | This paper |
| CD8a-(Htr6-CT)-BioID2-HA (Δ373-407) | CD8a(1-206)-BioID2-HA | Htr6 (326-440) + (Δ373-407) | XhoI-BamHI | This paper |
| CD8a-(Htr6-CT)-BioID2-HA (Δ373-389) | CD8a(1-206)-BioID2-HA | Htr6 (326-440) + (Δ373-389) | XhoI-BamHI | This paper |
| CD8a-(Htr6-CT)-BioID2-HA (Δ390-407) | CD8a(1-206)-BioID2-HA | Htr6 (326-440) + (Δ390-407) | XhoI-BamHI | This paper |
| CD8a-(Htr6-CT)-BioID2-HA (Δ408-440) | CD8a(1-206)-BioID2-HA | Htr6 (326-440) + (Δ408-440) | XhoI-BamHI | This paper |
| CD8a-(Htr6-CT)-BioID2-HA (Δ408-440+CTmut3) | CD8a(1-206)-BioID2-HA | Htr6 (326-440) + (Δ408-440) + (LLL398AAA) | XhoI-BamHI | This paper |
| CD8a-(Htr6-CT)-BioID2-HA (CTmut3) | CD8a(1-206)-BioID2-HA | Htr6 (326-440) + (LLL398AAA) | XhoI-BamHI | This paper |
| Sstr3-EGFP | pEGFP-N3 | Mouse Sstr3 (NP_033244.2) (428 aa) | XhoI-KpnI | Berbari et al. 2008a |
| Sstr3 (IC3mut1)-EGFP | pEGFP-N3 | Sstr3 + (A243F+Q247F+A251F+Q255F) | XhoI-KpnI | This paper |
| Sstr3 (IC3mut1+CTΔ1)-EGFP | pEGFP-N3 | Sstr3 (IC3mut1) + (Δ389-428) | XhoI-KpnI | This paper |
| Sstr3 (IC3mut1+CTΔ2)-EGFP | pEGFP-N3 | Sstr3 (IC3mut1) + (Δ355-388) | XhoI-KpnI | This paper |
| Sstr3 (IC3mut1+CTΔ3)-EGFP | pEGFP-N3 | Sstr3 (IC3mut1) + (Δ349-428) | XhoI-KpnI | This paper |
| Sstr3 (IC3mut1+CTΔ4)-EGFP | pEGFP-N3 | Sstr3 (IC3mut1) + (Δ335-348) | XhoI-KpnI | This paper |
| Sstr3 (IC3mut1+CTΔ5)-EGFP | pEGFP-N3 | Sstr3 (IC3mut1) + (Δ341-348) | XhoI-KpnI | This paper |
| Sstr3 (IC3mut1+CTΔ6)-EGFP | pEGFP-N3 | Sstr3 (IC3mut1) + (Δ335-340) | XhoI-KpnI | This paper |
| Sstr3 (IC3mut1+CTΔ7)-EGFP | pEGFP-N3 | Sstr3 (IC3mut1) + (Δ335-428) | XhoI-KpnI | This paper |
| Sstr3 (IC3mut1+CTmut1)-EGFP | pEGFP-N3 | Sstr3 (IC3mut1) + (LLRP337-340AARA) | XhoI-KpnI | This paper |
| Sstr3 (IC3mut1+CTmut2)-EGFP | pEGFP-N3 | Sstr3 (IC3mut1) + (FK329-330AA) | XhoI-KpnI | This paper |
| Sstr3 (CTmut1)-EGFP | pEGFP-N3 | Sstr3 (LLRP337-340AARA) | XhoI-KpnI | This paper |
| Sstr3 (IC3mut2+CTmut1)-EGFP | pEGFP-N3 | Sstr3 (CTmut1) + (VVK231-233AAA) | XhoI-KpnI | This paper |
| Sstr3 (IC3mut3+CTmut1)-EGFP | pEGFP-N3 | Sstr3 (CTmut1) + (VTR264-266AAA) | XhoI-KpnI | This paper |
| Sstr3 (IC3Δ1+CTmut1)-EGFP | pEGFP-N3 | Sstr3 (CTmut1) + (Δ234-242) | XhoI-KpnI | This paper |
| Sstr3 (IC3Δ2+CTmut1)-EGFP | pEGFP-N3 | Sstr3 (CTmut1) + (Δ243-247) | XhoI-KpnI | This paper |
| Sstr3 (IC3Δ3+CTmut1)-EGFP | pEGFP-N3 | Sstr3 (CTmut1) + (Δ248-255) | XhoI-KpnI | This paper |
| Sstr3 (IC3Δ4+CTmut1)-EGFP | pEGFP-N3 | Sstr3 (CTmut1) + (Δ243-255) | XhoI-KpnI | This paper |
| Sstr3 (IC3Δ5+CTmut1)-EGFP | pEGFP-N3 | Sstr3 (CTmut1) + (Δ256-263) | XhoI-KpnI | This paper |
| Sstr5-ECFP | pECFP-N1 | Mouse Sstr5 (NP_035555.1) (362 aa) | XhoI-KpnI | This paper |
| pG-LAP1-TULP3 | pG-LAP1 | Human TULP3 (NP_003315.2) (442 aa) | Gateway | Mukhopadhyay et al. 2010 |
| pG-LAP1-TULP3 (NTD) | pG-LAP1-TULP3 | TULP3 (aa 1-183) (replacing TULP3) | BstBI-XhoI | This paper |
| pG-LAP1-TULP3 (CTD) | pG-LAP1-TULP3 | TULP3 (aa 184-442) (replacing TULP3) | BstBI-XhoI | This paper |
| pcDNA3.1-TULP3-myc-his | pcDNA3.1-myc-his(-)C | Human TULP3 (NP_003315.2) (442 aa) | XhoI-BamHI | This paper |
| pcDNA3.1-TULP3(KR)-myc-his | pcDNA3.1-myc-his(-)C | pcDNA3.1-TULP3-myc-his + (K268A+R270A) | XhoI-BamHI | This paper |
| pEGFP-RABL2B | pEGFP-C1 | Human RABL2B (NP_001124393.1) (229 aa) | EcoRI-SalI | Dateyama et al. 2019 |
| pCMV5-Flag-RABL2B | pCMV5-Flag | Human RABL2B (NP_001336945.1) (235 aa) | EcoRI-SalI | Dateyama et al. 2019 |
| pcDNA3.1-B9D1-myc-his | pcDNA3.1-myc-his(-)C | Human B9D1 (NP_056496.1) (204 aa) | XhoI-KpnI | This paper |
